# Structural basis for Cas2/3 recruitment and nuclease-to-helicase DNA handover in type I-F CRISPR-Cas

**DOI:** 10.64898/2026.11.20.768062

**Authors:** Hyunsuh Kwon, Kazuki Ishihara, Takaaki Ogoshi, Christoph Gerle, Chai C. Gopalasingam, Hideki Shigematsu, Tomoyuki Numata

**Author notes:** To whom correspondence should be addressed. Laboratory of Biochemistry, Department of Bioscience & Biotechnology, Graduate School of Bioresource & Bioenvironmental Sciences, Kyushu University, 744 Motooka, Fukuoka 819-0395, Japan.

## Abstract

Type I CRISPR-Cas systems couple RNA-guided target recognition to DNA degradation by a Cas3 nuclease-helicase. However, how initial nicking of the displaced non-target strand (NTS) at the active site of the Cas3 HD nuclease domain is coordinated with its subsequent loading into the helicase module remains poorly understood. Here, we report cryo-electron microscopy structures of the canonical type I-F surveillance complex (Csy) from *Vibrio parahaemolyticus* in DNA-unbound and DNA-bound states, together with two Csy-DNA-Cas2/3 structures consistent with states before and after initial NTS cleavage. Target DNA binding elongates the Csy complex and repositions the Cas8 helical bundle domain, creating a Cas2/3 recruitment platform involving Cas8 and Cas5. Cas2/3 captures a flexible NTS segment as a bulge at its HD nuclease active site, where an open RecA1 gate loop permits substrate access. Comparison of the two Cas2/3-bound structures suggests that, following initial nicking, the PAM-proximal NTS product disengages from the Cas8 helical bundle domain and is redirected into the helicase channel. Together with biochemical analyses, these structures provide a mechanistic framework for understanding how initial nicking is coupled to loading of the cleaved NTS into the helicase channel in type I CRISPR-Cas systems.

## Introduction

Bacteria and archaea employ CRISPR-Cas systems as RNA-guided adaptive immune mechanisms against invading mobile genetic elements^1–4^. In these systems, effector complexes comprising one or more Cas proteins and a crR^5^NA recognize target nucleic acids through base pairing with the crRNA guide and direct their degradation^6–8^. CRISPR-Cas systems are broadly divided into two classes according to the architecture of their effector modules, with class 1 systems generally employing multisubunit effector complexes and class 2 systems using single multidomain effectors^9–11^. These two classes encompass seven types (I–VII), which are further divided into subtypes^11^. In canonical type I systems, the multisubunit CRISPR-associated complex for antiviral defense (Cascade) recognizes target DNA, whereas the Cas3 nuclease–helicase mediates its degradation^2,12–15^.

Target recognition by Cascade begins with binding to a protospacer-adjacent motif (PAM), followed by base pairing between the crRNA guide and the complementary target strand (TS) of the DNA^13,14,16,17^. This generates an R-loop in which the non-target strand (NTS) is displaced as single-stranded DNA^18–21^. Cas3 contains an HD nuclease domain and a superfamily 2 (SF2) helicase module comprising two RecA-like domains, RecA1 and RecA2^12,22,23^. The HD domain introduces an initial nick into the NTS, whereas the helicase drives ATP-dependent 3′-to-5′ translocation and delivers the NTS to the HD active site for processive degradation^15,23^. Target recognition controls Cas3 function through distinct mechanisms. In characterized type I-C, I-E, and I-F systems, R-loop formation enables Cas3 recruitment^5,15,24–26^, whereas in the *Pyrococcus furiosus* type I-A system, Cas3 is constitutively associated with Cascade and is activated allosterically upon full R-loop formation^27^. In addition to Cas3 recruitment, access of the NTS to the HD nuclease active site is subject to regulation. Structures of the *Pseudomonas aeruginosa* type I-F Cas1-Cas2/3 complex, in which Cas2/3 is a Cas2-Cas3 fusion protein^10,25,28^, revealed a RecA1-derived loop that occludes the HD active site, suggesting that relief of this steric barrier contributes to nuclease activation^29^.

The canonical type I-F1 Cascade, hereafter referred to as the Csy complex, comprises four distinct Cas proteins, Cas5, Cas6, Cas7, and Cas8, assembled around a crRNA^30,31^. Biochemical and structural studies of the *P. aeruginosa* Csy complex (PaeCsy) revealed how PaeCsy recognizes a 5′-CC-3′ PAM and undergoes extensive structural rearrangements upon target DNA binding^5,21^. Full R-loop formation induces elongation of the Cas7 backbone and an approximately 180° rotation of the Cas8 helical bundle (HB) domain, exposing a nuclease recruitment helix implicated in Cas2/3 binding^5^. Alternative recruitment mechanisms have been identified in noncanonical type I-F systems. In ICP1, a bacteriophage infecting *Vibrio cholerae*, Cas8 lacks the HB domain, and Cas1 bridges the DNA-bound Csy complex and Cas2/3^32,33^. Type I-F2 systems lack Cas8 altogether and instead recruit Cas2/3 through contacts with Cas7 and Cas5^34–36^. Structures of these noncanonical complexes reveal the diversity of recruitment mechanisms but do not establish how Cas2/3 engages the canonical Csy complex through the Cas8 HB domain.

Structural studies have also illuminated how Cas3 processes the displaced NTS. Structures of type I-E Cascade-DNA-Cas3 complexes revealed a helicase-bypassing route for initial nicking and subsequent relocation of the cleaved NTS toward the helicase entrance^15,17^. A type I-C Cascade-DNA-Cas3 structure resolved an NTS bulge captured at the HD active site^26,37^, whereas a type I-A structure showed the nicked PAM-proximal NTS segment threaded through the helicase module toward the nuclease domain^27^. In noncanonical type I-F systems, structures of ICP1 and type I-F2 Csy complexes bound to DNA and Cas2/3 revealed NTS engagement by the helicase module^33,36^, but did not capture the NTS positioned for initial nicking at the HD active site. These studies provide structural insights into distinct stages of NTS processing. However, it remains unclear how the NTS trajectory and its interactions with surrounding proteins change during transfer from the HD active site into the helicase channel. In particular, in canonical type I-F systems, neither NTS capture at the HD active site nor accommodation of the cleavage product within the helicase channel has been structurally characterized. Comparison of structures representing these two stages is therefore essential for establishing how changes in the NTS trajectory and its interactions with Csy and Cas2/3 couple initial nicking to loading of the cleaved NTS into the helicase channel.

Here, we determined cryo-electron microscopy (cryo-EM) structures of the canonical type I-F Csy complex from *Vibrio parahaemolyticus* in the absence and presence of target DNA, together with two distinct states of the Csy-DNA-Cas2/3 complex. These structures directly define the interface between the Cas8 HB domain and Cas2/3, along with additional contacts between Cas5 and Cas2/3, revealing how target DNA-induced Csy rearrangements create a platform for Cas2/3 recruitment. The two Cas2/3-bound structures are consistent with states before and after initial NTS cleavage. Their comparison reveals distinct NTS trajectories and associated changes in contacts with the Cas8 HB domain and Cas5, providing a structural basis for the transition from NTS bulge capture at the HD active site to accommodation of the PAM-proximal cleavage product within the helicase channel. Together with biochemical analyses, these structures provide a framework linking target recognition, Cas2/3 recruitment, initial nicking, and cleavage-product handover in canonical type I-F CRISPR-Cas interference.

## Results

### Structural rearrangements of the *V. parahaemolyticus* Csy complex upon target DNA binding

To evaluate the functionality of the type I-F CRISPR-Cas system identified in *V. parahaemolyticus* 3259 (Fig. 1a), we first performed an *in vivo* plasmid interference assay. For this experiment, we prepared a target plasmid containing a 5′-CC-3′ PAM and a protospacer complementary to the first spacer of CRISPR locus. The experiment revealed that the type I-F CRISPR-Cas is functional and capable of mediating plasmid interference in this heterologous *E. coli* expression system (Fig. 1b). We next reconstituted the *V. parahaemolyticus* Csy complex (VpCsy) in *E. coli* cells by co-expressing the Cas proteins and crRNA, followed by purification of the assembled complex (Fig. 1c). *V. parahaemolyticus* Cas2/3 (VpCas2/3) was also produced in *E. coli* cells and purified. To examine target DNA cleavage by VpCsy and VpCas2/3, a double-stranded target DNA containing a 5′-Cy5-labeled NTS was incubated in the presence or absence of VpCsy and VpCas2/3. The results showed that VpCsy- and VpCas2/3-mediated cleavage initially produced a fragment migrating close to the 50-nt marker (Fig. 1d). Over the time course, this product was progressively accumulated and concomitantly degraded to a final fragment of approximately 10-nt (Fig. 1d), consistent with processive DNA degradation by the Cas2/3 nuclease/helicase following initial nicking.

**Figure 1.**
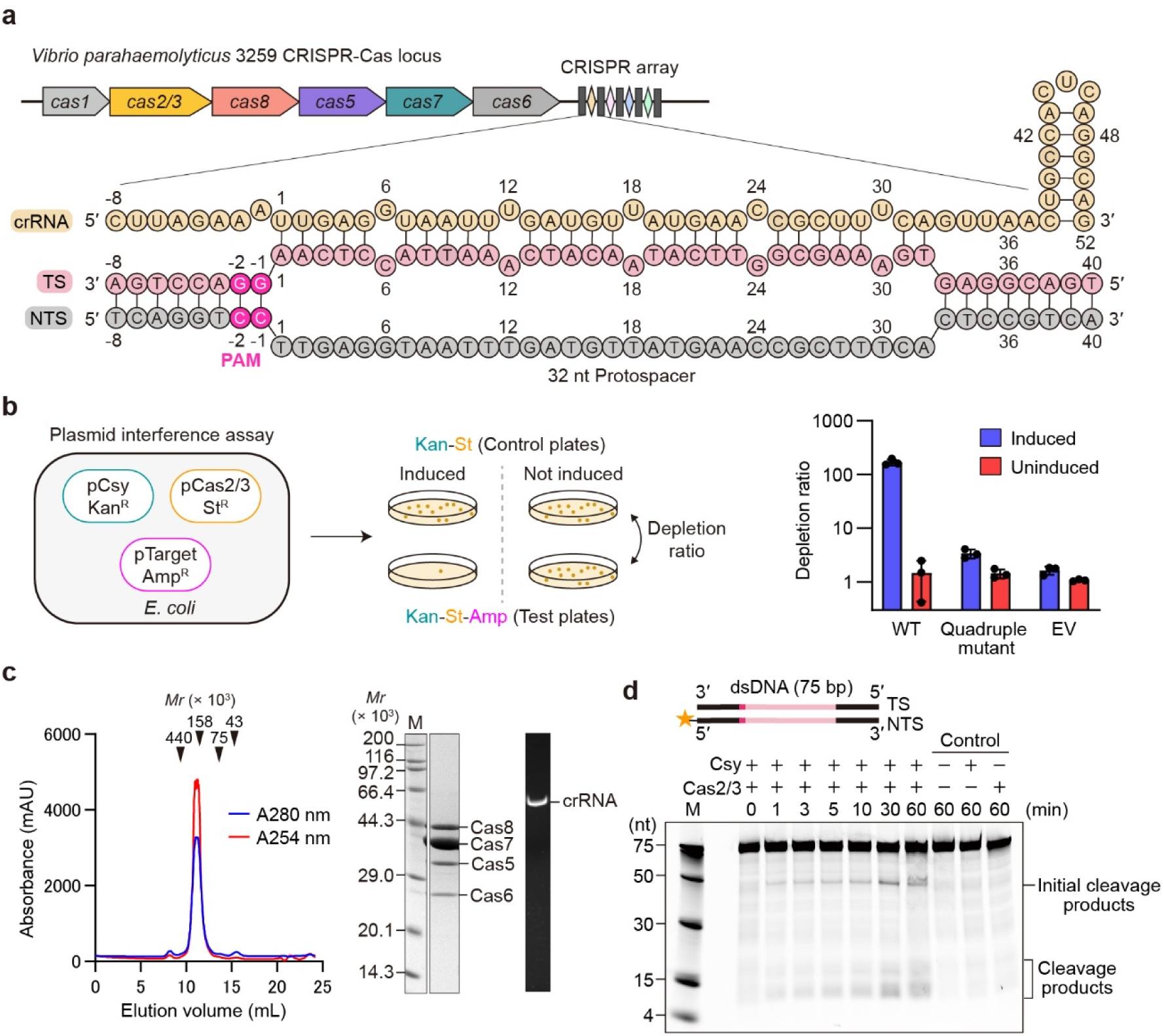
Plasmid interference and DNA cleavage activities of the *V. parahaemolyticus* type I-F CRISPR-Cas system. **(a)** Schematic representation of the type I-F CRISPR-Cas locus in *V. parahaemolyticus* 3259 (top) and the crRNA-target DNA R-loop (bottom). The crRNA derived from the first spacer of the CRISPR array is shown in wheat, the target strand (TS) in salmon, and the non-target strand (NTS) in gray. The protospacer-adjacent motif (PAM) is highlighted in hot pink. The 32-nt protospacer and its flanking sequences are shown. **(b)** *In vivo* plasmid interference assay. Left, schematic overview of the assay. *E. coli* cells harboring pCsy (kanamycin resistance), pCas2/3 (streptomycin resistance), and pTarget (ampicillin resistance) were plated on LB agar containing kanamycin and streptomycin (control plates) or kanamycin, streptomycin, and ampicillin (test plates), with or without IPTG. Right, depletion ratios for wild-type (WT) VpCsy, the Cas8 helical bundle (HB) quadruple mutant, and the empty-vector control (EV) under induced and uninduced conditions. The depletion ratio was calculated by dividing the number of colonies recovered on control plates by that recovered on test plates. Bars represent the mean ± SD of three independent experiments. Individual data points are shown as black circles. The Cas8 HB quadruple mutant contains the Y313A, N315A, I316A, and R319A substitutions, targeting residues involved in TS contacts described below (Fig. 2e). **(c)** Purification of the VpCsy complex. Left, size-exclusion chromatography profile obtained using a Superdex 200 Increase 10/300 GL column. Arrowheads indicate the elution positions of molecular mass standards: ferritin (440 kDa), aldolase (158 kDa), conalbumin (75 kDa), and ovalbumin (44 kDa). Middle and right, SDS-PAGE and denaturing PAGE analyses of the purified complex, showing its protein subunits and crRNA, respectively. M, molecular mass markers. **(d)** Time course of target DNA cleavage by purified VpCsy and VpCas2/3. A schematic of the 75-bp dsDNA substrate is shown above the gel. The NTS was labeled at its 5′ end with Cy5 (orange star). Reactions were analyzed at the indicated time points. Control reactions lacking both proteins or either protein were incubated for 60 min. The positions of the initial cleavage products and shorter cleavage products are indicated. M, DNA size markers. The gel image is representative of three independent experiments.

To characterize the structural features of VpCsy, we prepared VpCsy alone and in complex with target DNA (Fig. S1) and determined their structures by cryo-EM single-particle analysis at resolutions of 2.10 Å and 1.98 Å, respectively (Figs. 2a, 2b, S2, and S3; Table S1). Similar to Csy complexes from other species, VpCsy comprises six Cas7 subunits, one copy each of Cas5, Cas6, and Cas8, and a single crRNA molecule, forming an overall spring washer-like architecture (Fig. 2a). Cas5 and Cas7.6 bind the 5′ tag of the crRNA, Cas7.1–Cas7.6 form the backbone of the complex, Cas6 recognizes the 3′ hairpin of the crRNA, and Cas8 bridges the top and middle regions of the complex through its interaction with Cas5 (Fig. 2a). Hereafter, the structures of VpCsy alone and VpCsy with target DNA are referred to State I and State II, respectively.

**Figure 2.**
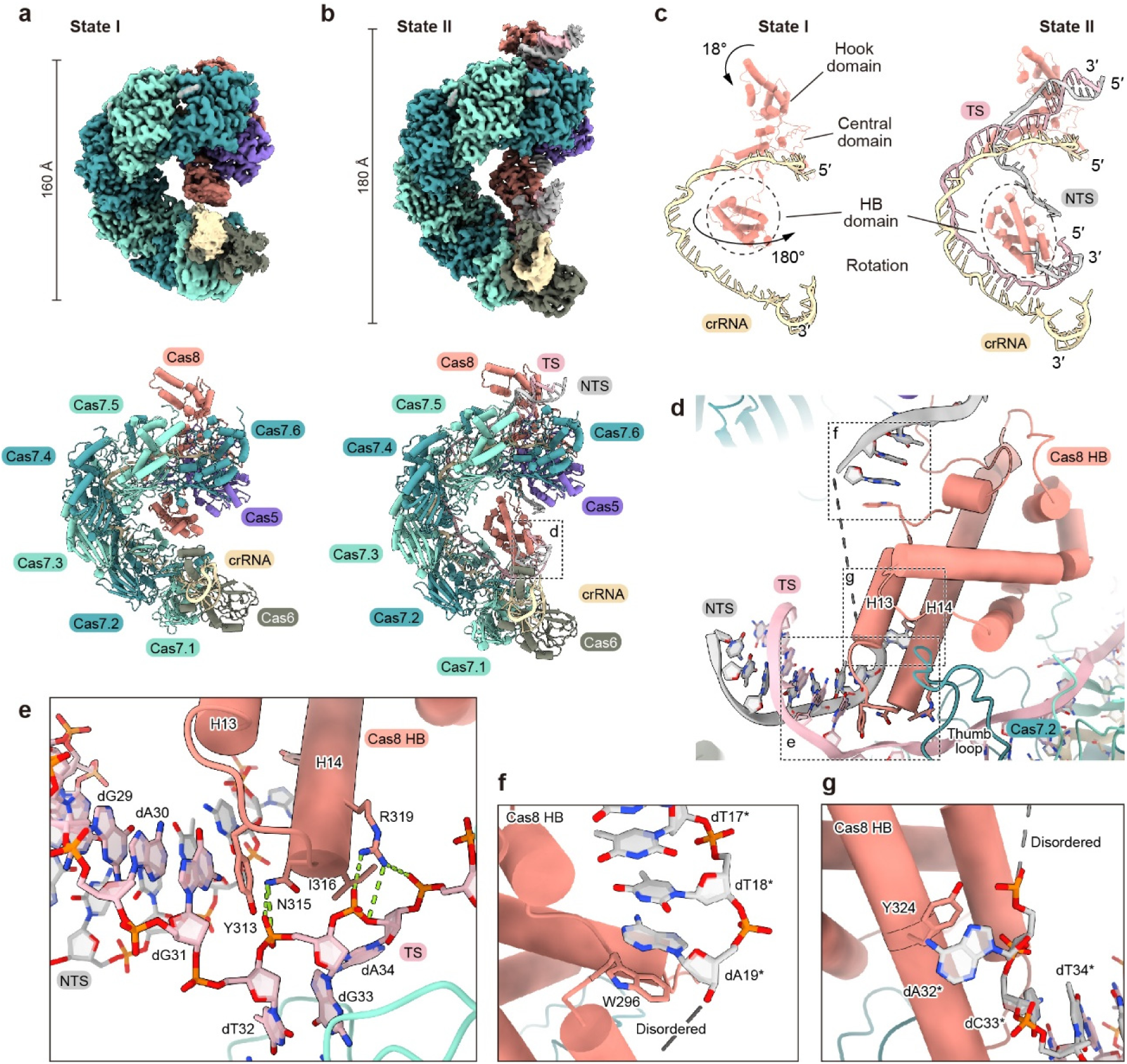
Cryo-EM structures of the VpCsy complex in target DNA-unbound and -bound states. **(a, b)** Overall structures of the target DNA-unbound (State I; **a**) and target DNA-bound (State II; **b**) VpCsy complexes. Cryo-EM maps (top) and cartoon representations of the atomic models (bottom) are colored according to the labeled components. Target DNA binding increases the overall length of the complex from approximately 160 to 180 Å. The dashed box in **b** indicates the region enlarged in **d**. **(c)** Comparison of Cas8 and the associated nucleic acids in States I and II, highlighting the rotation of the Cas8 HB domain upon target DNA binding. The hook, central, and HB domains of Cas8 are indicated. Cas8 is shown in salmon, crRNA in wheat, the TS in pink, and the NTS in gray. **(d)** Close-up view of the Cas8 HB domain and target DNA in State II, showing contacts of the loop connecting helices H13 and H14 with the PAM-distal DNA duplex and additional contacts of the HB domain with the displaced NTS. Dashed boxes indicate the regions enlarged in **e–g**. Gray dashed lines indicate disordered NTS segments. **(e)** Close-up view of interactions of the H13–H14 loop and adjacent residues with the TS near the PAM-distal end of the R-loop. Hydrogen bonds are indicated by light green dashed lines. **(f, g)** Close-up views of contacts of Trp296 (**f**) and Tyr324 (**g**) with NTS nucleotides dT18* and dA32*, respectively.

Cas8 consists of three major domains, the hook domain, the central domain, and the helical bundle (HB) domain (Fig. S4a)^21,31^. State II structure reveals 5′-CC-3′ PAM recognition mechanism of VpCsy complex where the side chains of Gln253 and Asn254 of Cas8 form base-specific hydrogen bonds with dC(-1)* and dG(-1), respectively (asterisk indicates nucleotide on the NTS) (Fig. S4b). The side chain of Asn114 also forms a hydrogen bond with exocyclic amino group of dG(-2), a nucleotide complementary to dC(-2)* (Fig. S4b). The PAM binding pose is similar to that reported in the structure of PaeCsy complex, consistent with amino acid sequence similarity around PAM recognition site (Fig. S4c). Structural comparison of the DNA-unbound and DNA-bound VpCsy complexes reveals substantial conformational rearrangements upon target DNA binding. The Cas8 hook domain moves approximately 18° toward the target DNA (Fig. 2c). Consistent with previous structures of the PaeCsy complex^5^, the Cas8 HB domain undergoes an approximately 180° rotation, thereby exposing a helix (residues 376–396) corresponding to the previously-reported Cas2/3 nuclease recruitment helix of PaeCas8 (Figs. 2a–2c). These rearrangements are accompanied by an approximately 20 Å extension of the VpCsy complex (Figs. 2a–2c).

In the DNA-unbound VpCsy structure, the Cas8 HB domain does not appear to interact tightly with the Cas7 backbone. By contrast, in the DNA-bound structure, the Cas8 HB domain interacts with the thumb loop of Cas7.2, a characteristic structural element of Cas7 family proteins (Fig. 2d)^19,38^. The Cas8 HB domain also contacts both the TS and the NTS at the PAM-distal end of the R-loop, beyond which the target DNA remains double-stranded (Fig. 2d). A loop between helices H13 and H14 (residues 295– 346) of the Cas8 HB domain contacts the TS (Fig. 2d). Tyr313 and Ile316 form stacking interactions with dG31 and dA34, respectively, while Asn315 and Arg319 form hydrogen bonds with the phosphate backbone (Fig. 2e). Notably, Tyr313 is positioned near the junction between the R-loop and the adjoining dsDNA (Fig. 2e). To assess the functional contribution of these Cas8 HB-DNA interactions, we generated a quadruple mutant (Y313A/N315A/I316A/R319A) and examined its activity using the plasmid interference assay. Colony-forming units obtained with the quadruple mutant was comparable to that obtained with the empty-vector control (Fig. 1b), indicating a marked reduction in target plasmid interference activity.

Whereas continuous cryo-EM density for the TS was observed from dG14 to dT62, nucleotides 20–31 of the NTS were disordered and could not be modeled. In contrast, dA19* and dA32* are well ordered and formed stacking interactions with Trp296 and Tyr324, respectively (Figs. 2f and 2g). These observations indicate that part of the NTS is stabilized through interactions with the Cas8 HB domain, whereas the adjacent NTS region remains flexible. Together, these structural and functional observations support the contribution of the Cas8 HB-DNA interactions to CRISPR-Cas interference.

### Cryo-EM structures of VpCsy-DNA-Cas2/3 complex

To elucidate the structural basis for VpCas2/3 recruitment and activation after target DNA binding, we prepared VpCsy complexed with both target DNA and VpCas2/3 (Fig. S5). Then, we determined its cryo-EM structures by single-particle analysis at resolutions of 2.26 Å and 2.58 Å in two distinct conformations (referred to as State III and State IV) from a single dataset, respectively (Figs. 3a, 3b, and S6; Table S1). In both structures, the N-terminal Cas2 moiety of VpCas2/3 was disordered, likely owing to the flexible linker connecting the Cas2 and Cas3 moieties. In contrast, the C-terminal Cas3 moiety was well resolved (Figs. 3a and 3b). Cas3 comprises an HD nuclease domain, a superfamily 2 (SF2) helicase composed of two RecA-like domains (RecA1 and RecA2), a linker domain, and a C-terminal domain (CTD) (Fig. 3c).

**Figure 3.**
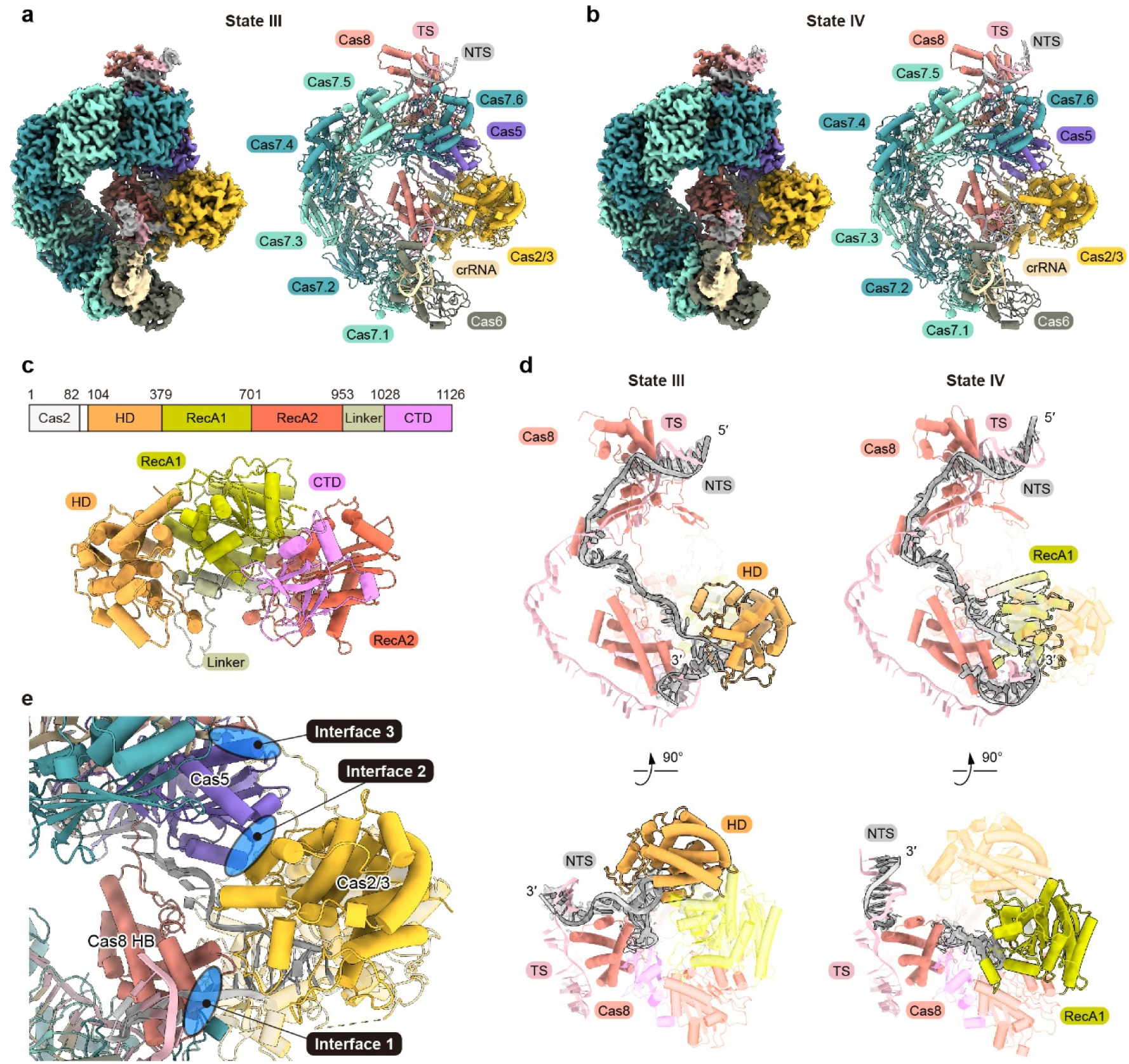
Cryo-EM structures of the VpCsy-dsDNA-VpCas2/3 complex in two distinct states. **(a, b)** Overall structures of the complex in State III (**a**) and State IV (**b**). Cryo-EM maps (left) and cartoon representations of the atomic models (right) are colored according to the labeled components. **(c)** Domain organization and structure of VpCas2/3. Top, schematic representation of the domain organization, with residue numbers indicating domain boundaries. Bottom, cartoon representations of VpCas2/3 in State III colored according to the schematic. The Cas2 domain is not resolved in the cryo-EM map. **(d)** Comparison of NTS conformations in State III (left) and State IV (right), shown in two orientations related by a 90° rotation. In State III, the HD domain of VpCas2/3 is highlighted, with all other domains shown as semitransparent cartoons. In State IV, the RecA1 domain is highlighted, with all other domains shown as semitransparent cartoons. Cas8 and the NTS are colored salmon and gray, respectively. VpCas2/3 domains are colored as in **c**. Dashed lines indicate disordered NTS segments. **(e)** Overview of the three interfaces between VpCsy and VpCas2/3. Interface 1 involves the Cas8 HB domain, interface 2 involves Cas5, and interface 3 involves both Cas5 and Cas8. The interfaces are highlighted in blue.

Interestingly, comparison of the State III and State IV structures reveals marked difference in the paths of the NTS (Fig. 3d). In the State III structure, nearly continuous density for the NTS extends toward the nuclease active site of the HD domain of VpCas2/3 (Fig. 3d). In the State IV structure, the NTS density is discontinuous, and the PAM-proximal region of the NTS extends toward the RecA-like helicase domains (Fig. 3d). The PAM-distal portion of the NTS (nucleotides 31–32) in the State IV structure remains separated from this region and, further downstream, pairs with the TS to form dsDNA (Fig. 3d). The two resolved NTS segments in the State IV structure are too far apart to be connected as a continuous single strand (see below). Therefore, these two structures likely represent pre- and post-NTS-cleavage states, respectively. The overall structures of VpCsy in the pre- and post-NTS-cleavage states are similar, whereas the positions of VpCas2/3 differ slightly between the two states (see below), likely reflecting the presence of either the intact NTS or the NTS cleavage products.

The structures show that VpCas2/3 binds VpCsy predominantly through three interfaces (Fig. 3e). The first interface is formed between the Cas8 HB domain and the Cas2/3 CTD, whereas the other two interfaces involve Cas5 and either the Cas2/3 HD domain or linker domain. At the first interface, Gln1056, Thr1057, Asn1059, and Gln1111 in the Cas2/3 CTD contact Arg387, Asp405, and Leu408 on the nuclease recruitment helix (residues 376–396) of the Cas8 HB domain, predominantly through hydrogen bonds (Fig. S7a). VpCas2/3 additionally interacts with Cas5 at the second interface, where Asn232, Arg234, Asp235, and Lys236 in the Cas2/3 HD domain contact Arg2, Asp114, Glu117, Leu149, and Glu157 of Cas5, primarily through electrostatic interactions (Fig. S7b). The third interface is mediated predominantly by interactions between Cas5 and the anchor loop (residues 970–998) of Cas2/3 linker domain (Fig. S7c). This interface also includes interactions between this anchor loop and a loop in the Cas8 central domain (residues 181–196), hereafter termed the nuclease recruitment loop (Fig. S7c). Within this interface, the negatively charged residues Glu188 of Cas8 and Glu39, Glu108, and Asp270 of Cas5 engage a cluster of basic residues on Cas2/3, including Arg984, Arg989, and Arg990 (Figs. 3e and S7c). Furthermore, Phe985 of Cas2/3 is inserted into a surface groove formed jointly by Cas8 and Cas5 and forms a hydrophobic core together with their hydrophobic residues (Fig. S7c), suggesting that this interaction contributes to stabilization of the third interface.

### Target DNA binding enables Cas2/3 recruitment through Csy rearrangement

Structural comparison of State II and State III revealed that the overall structures of the Csy complex, including the relative position of the Cas8 HB domain, are well superimposed (Fig. S8). Thus, Cas2/3 binding does not induce a substantial additional rearrangement of the Csy complex after target DNA binding. Together with the structural rearrangements observed upon target DNA binding described above (Figs. 2a–2c), these observations suggest that the target DNA-bound Csy complex adopts a preorganized architecture for Cas2/3 recruitment.

VpCas2/3 interacts with VpCsy mainly through the Cas8 HB domain and Cas5, as described above (Figs. 3e and S7). However, the position of Cas5 is similar between State I and State II, whereas the Cas8 HB domain undergoes a large rearrangement upon the target DNA binding (Figs. 2a–2c). These observations suggest that the ability of Csy to recruit Cas2/3 is primarily controlled by target DNA binding and the subsequent structural rearrangement of the Cas8 HB domain.

Furthermore, superimposition of State I and State III showed that Cas2/3 would sterically clash with the Cas7.1 subunit if it were positioned on the DNA-unbound State I structure (Fig. 4). This steric clash arises from the more compact architecture of Csy in the absence of target DNA and is relieved by extension of the Csy complex upon target DNA binding (Fig. 4). Thus, target DNA binding converts VpCsy into a conformation that can accommodate VpCas2/3.

**Figure 4.**
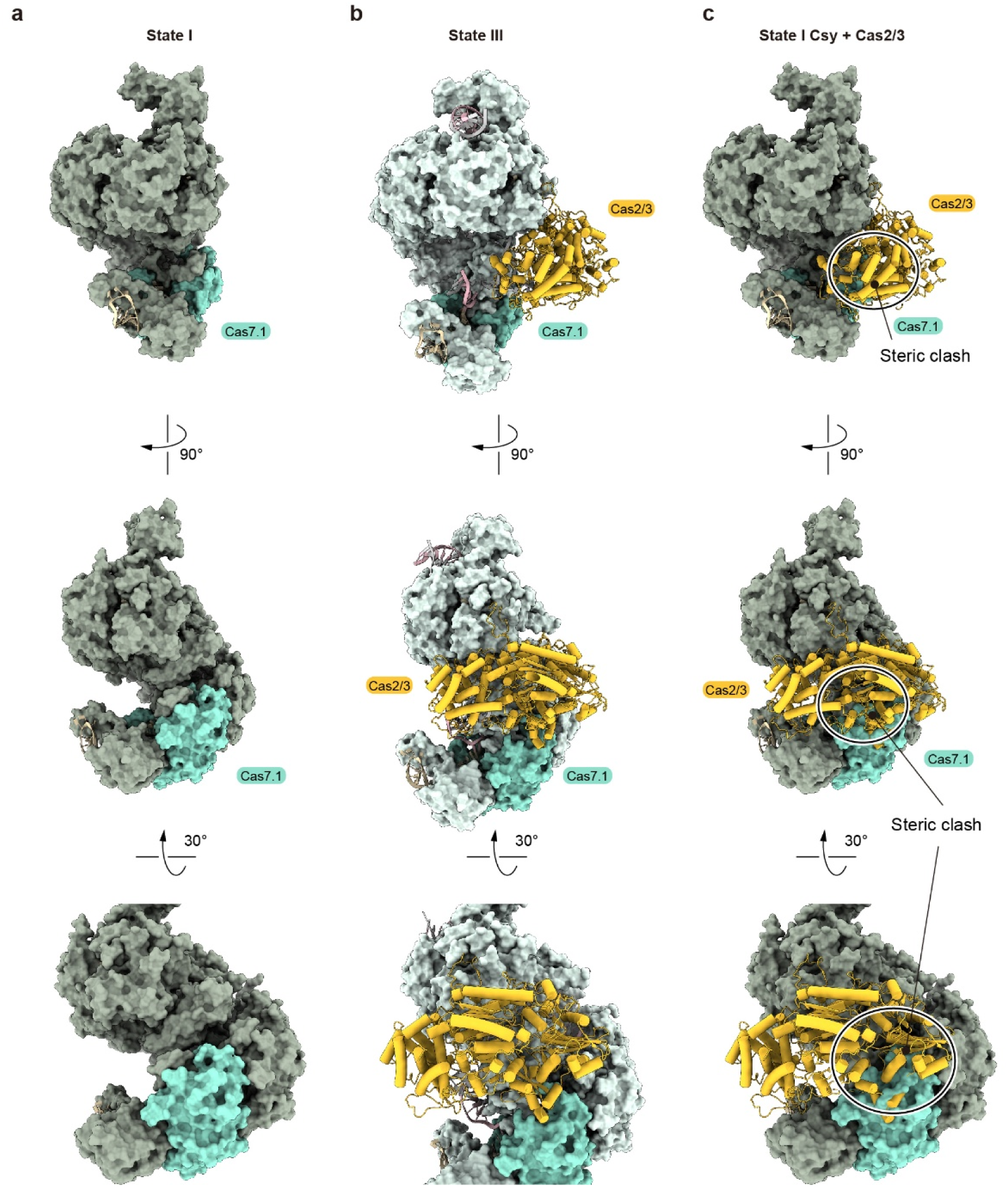
Accommodation of Cas2/3 to the target DNA bound-Csy complex. **(a)** Surface representation of the target DNA-unbound Csy complex (State I). Cas7.1 is colored light green, and the remaining protein subunits are colored dark gray. **(b)** Structure of the Csy-dsDNA-Cas2/3 complex (State III). Csy protein subunits are shown as surfaces, with Cas7.1 colored light green and the remaining subunits colored light gray. Cas2/3 is shown as a yellow cartoon. **(c)** Model of Cas2/3 positioned on the target DNA-unbound Csy complex (State I) based on structural superimposition with State III. Circles indicate steric clashes between Cas7.1 and Cas2/3. The Csy complex is colored as in **a**, and Cas2/3 is shown as a yellow cartoon. In **a–c**, the top and middle views show the overall structures in two orientations related by a 90° rotation. The bottom views show close-ups of the Cas7.1 region after an additional 30° rotation relative to the middle views.

### NTS positioning in the Cas2/3 HD active site and Cas2/3 activation

In the State III structure, nucleotides dT20*–dA32* of the NTS form a bulge that is accommodated within the Cas2/3 HD nuclease domain (Figs. 5a and 5b). The corresponding segment is disordered in the state II structure, suggesting that Cas2/3 recruitment stabilizes this flexible NTS segment in a bulged conformation within the HD domain. Importantly, the phosphodiester bond between dC24* and dC25* is sharply kinked at the HD active site, where Asp124, Lys127, His219, Asp324, and His360 are positioned to contact the phosphate group (Fig. 5b). Two divalent metal ions are coordinated by His123, Asp124, His151, His218, His219, Asp324, and His360, with Asp124 coordinating both divalent cations (Fig. 5b). The two non-bridging oxygen atoms of the phosphate group each coordinate one metal ion (Fig. 5b). These active-site residues are conserved among Cas2/3 family proteins (Fig. S9a), and a similar active-site architecture is observed in PaeCas2/3 (Fig. S9b)^39^. These observations suggest that the phosphodiester bond between dC24* and dC25* is positioned for cleavage by the Cas2/3 HD domain, consistent with the production of an approximately 50-nt fragment when a Cy5-labeled 75-bp dsDNA was used as the substrate (Fig. 1d).

**Figure 5.**
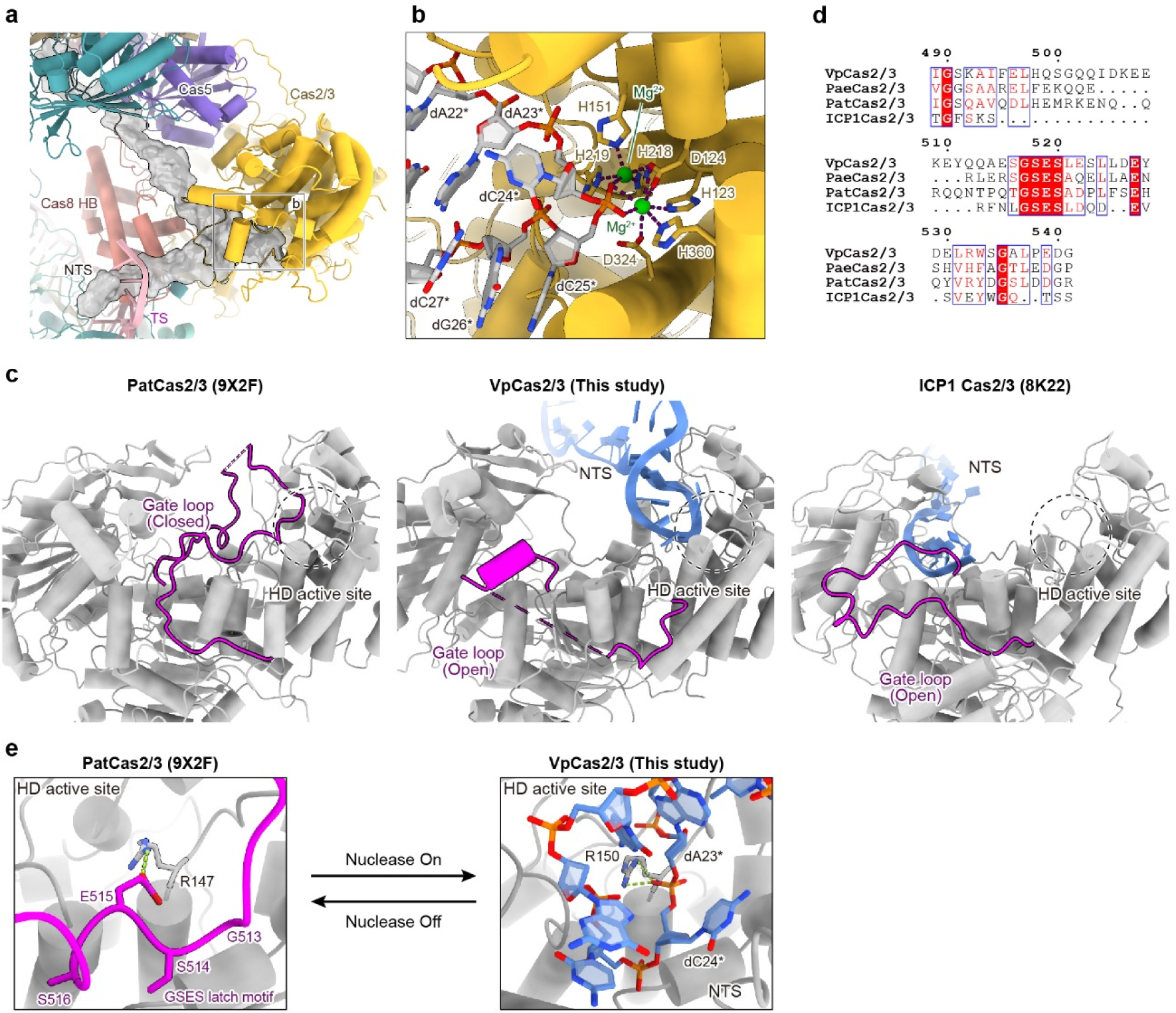
Positioning of the NTS bulge in Cas2/3 HD active site in State III. **(a)** Structure of the NTS bulge within the Csy-dsDNA-Cas2/3 complex. The NTS is shown as a gray cartoon surrounded by a semitransparent gray surface. The remaining components are shown as cartoons. Cas5, Cas7.6, Cas2/3, the Cas8 HB domain, and the TS are colored purple, teal, yellow, salmon, and pink, respectively. The box indicates the region enlarged in **b**. **(b)** Close-up view of the Cas2/3 HD active site. The NTS segment and selected amino acid residues are shown as sticks. Two divalent metal ions, modeled as Mg²⁺, are shown as green spheres, with the corresponding cryo-EM densities overlaid as semitransparent surfaces. **(c)** Comparison of RecA1 gate loop conformations in PatCas2/3 (PDB ID: 9X2F), VpCas2/3 (State III; this study), and ICP1 Cas2/3 (PDB ID: 8K22). The gate loop is shown in magenta, and the remaining protein regions are shown in gray. PatCas2/3 adopts a closed gate loop conformation, whereas VpCas2/3 and ICP1 Cas2/3 adopt open conformations. Disordered gate loop segments are indicated by dashed lines. The NTS is shown as a blue cartoon, and dashed circles indicate the HD active site. **(d)** Amino acid sequence alignment of the RecA1 gate loop regions of VpCas2/3, PaeCas2/3, PatCas2/3, and ICP1 Cas2/3. Residue numbers of VpCas2/3 are indicated above the alignment. **(e)** Comparison of interactions involving the conserved arginine residue near the HD active site in PatCas2/3 (PDB ID: 9X2F; left) and VpCas2/3 in State III (right). In PatCas2/3, Arg147 forms hydrogen bonds with Glu515 of the GSES latch motif in the closed RecA1 gate loop. In VpCas2/3, the corresponding Arg150 contacts the phosphate groups of NTS nucleotides dA23* and dC24*. Selected residues and nucleotides are shown as sticks. The gate loop is colored magenta, the remaining protein regions gray, and the NTS blue. Putative hydrogen bonds are indicated by green dashed lines. Arrows illustrate the proposed transition between inactive and active nuclease states.

We next examined conformational differences of Cas2/3 between the Csy-bound and Csy-unbound states. Since no structure of VpCas2/3 alone is available, we compared VpCas2/3 in state III with the structure of *Pectobacterium atrosepticum* Cas2/3 (PatCas2/3). Superimposition of the PatCas2/3 structure onto VpCas2/3 in State III revealed a prominent conformational difference in the RecA1 gate loop (residues 489– 541 of VpCas2/3), a loop extending from the RecA1 domain that is proposed to regulate substrate access to the HD active site (Fig. 5c)^29^. In PatCas2/3, the RecA1 gate loop is well ordered and covers the HD active site (Fig. 5c). The GSES latch motif, conserved among type I-F Cas2/3 proteins, forms hydrogen bonds with the HD domain, stabilizing the closed conformation of the gate loop (Fig. 5d and 5e). This conformation has been proposed to maintain Cas2/3 in an inactive state by occluding the substrate channel and preventing ssDNA from accessing the HD nuclease domain^29,40^. By contrast, the corresponding region in VpCas2/3 in state III extends away from the HD active site, and residues 501–530 are disordered, resulting in an open HD active site (Fig. 5c and 5d). This open architecture allows the NTS bulge to be accommodated within the HD domain (Figs. 5a and 5c). In State III, the conserved Arg150 of VpCas2/3 forms hydrogen bonds with the phosphate groups of NTS nucleotides dA23* and dC24*, whereas the corresponding arginine in PatCas2/3 interacts with the conserved glutamate of the GSES latch motif (Fig. 5e).

Comparison of the closed PatCas2/3 structure with VpCas2/3 in State III therefore suggests that Csy recruitment is accompanied by remodeling of the RecA1 gate loop, relieving the steric block and allowing the NTS to access the HD active site. The Csy complex of ICP1, a phage that infects *Vibrio cholerae*, in noncanonical: it lacks the Cas8 HB domain and recruits Cas2/3 through a Cas1-mediated mechanism distinct from that of canonical type I-F Csy complexes such as VpCsy. Despite this difference, the RecA1 gate loop in the ICP1 Csy-dsDNA-Cas1-Cas2/3 complex is similarly displaced away from the HD active site (Fig. 5c)^33^. However, the resolved NTS in the ICP1 complex occupies the helicase channel without extending into the HD active site, whereas in VpCas2/3 State III, it forms a bulge accommodated at the HD active site (Fig. 5c).

Together, these observations suggest that Cas2/3 recruitment couples opening of the RecA1 gate with stabilization of the flexible NTS segment at the HD active site, thereby positioning the NTS for initial cleavage.

### Handover of the nicked NTS product to the RecA1 and RecA2 helicase domains

In State IV, nucleotides dT18*–dC24* of the NTS are accommodated within a basic channel formed by the RecA1 and RecA2 domains and the CTD of VpCas2/3 (Fig. 6a). The sugar-phosphate backbone of the NTS contacts Arg795, Arg850, and Lys895 in RecA2 (Fig. 6a). Several hydrophilic residues, including Ser788 and Thr843, as well as hydrophobic residues, including Leu463, Ile759, Pro844, and Val853, also contribute to NTS binding (Fig. 6a). Notably, a loop connecting the linker domain and the CTD (residues 1020–1029) arches over nucleotides dA23* and dC24* of the NTS, likely clamping this NTS segment within the cleft (Fig. 6a).

**Figure 6.**
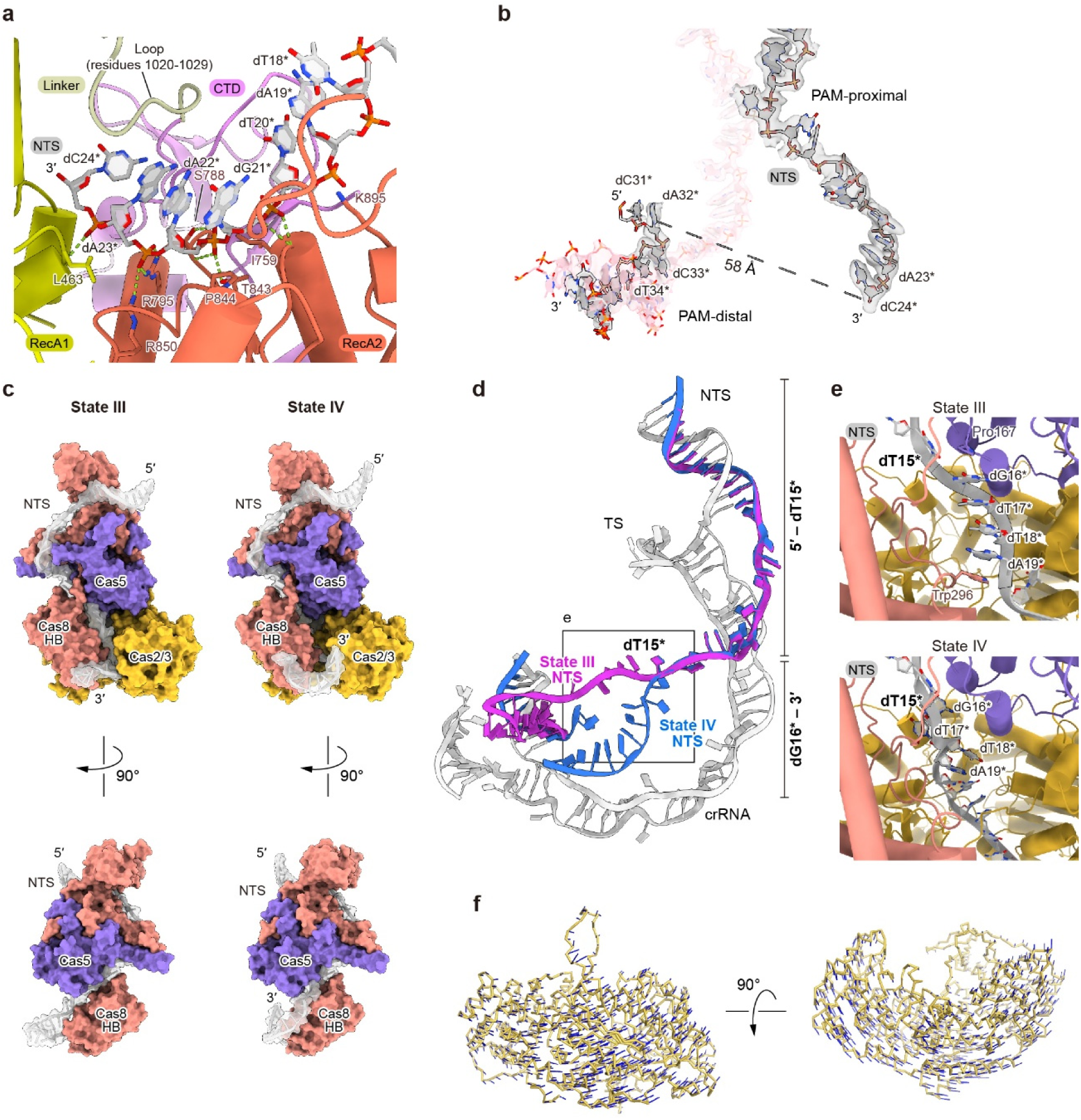
NTS accommodation in the Cas2/3 helicase channel and structural rearrangements between States III and IV. **(a)** Interactions between the NTS and Cas2/3. The NTS and selected interacting amino acid residues are shown as sticks. Putative hydrogen bonds are indicated by green dashed lines. The RecA1, RecA2, linker domain, and CTD are colored yellow, orange, ivory, and purple, respectively, and the NTS is colored light gray. The loop connecting the linker domain and CTD (residues 1020–1029) is indicated. **(b)** Spatial separation of the two resolved NTS segments in State IV. The distance between the O3′ atom of dC24* and the phosphorus atom of dC31* is 54 Å, as indicated by the dashed line. The NTS and TS are shown as sticks, with the corresponding cryo-EM densities overlaid as semitransparent surfaces. The PAM-proximal and PAM-distal regions are labeled. **(c)** Comparison of NTS trajectories in State III (left) and State IV (right), shown in two orientations related by a 90° rotation. The NTS is colored light gray and shown with semitransparent surfaces. Cas5, Cas8, and Cas2/3 are shown as purple, salmon, and yellow surfaces, respectively. Cas2/3 is omitted from the bottom views to reveal the NTS trajectory. **(d)** Superimposition of the R-loop structures in States III and IV. The NTS is colored magenta in State III and blue in State IV, whereas the TS and crRNA are colored light gray. The point at which the two NTS trajectories diverge, at dT15*, is labeled. The box indicates the region shown in **e**. **(e)** Close-up views of NTS trajectories between Cas5, Cas8, and Cas2/3 in State III (top) and State IV (bottom). The NTS is shown as a gray cartoon, with selected nucleotides labeled. Cas5 Pro167 and Cas8 Trp296, which participate in a continuous stacking interaction with dT15*–dT18* in State III, are shown as sticks. **(f)** Rotational shift of Cas2/3 from State III to State IV, shown in two orientations related by a 90° rotation. Cas2/3 in State III is shown as a yellow Cα trace. Blue lines connect corresponding Cα atoms in States III and IV, indicating their positional displacements. For clarity, the Cα trace of Cas2/3 in State IV is omitted.

Density for the PAM-proximal NTS segment is well defined through dC24*, with much weaker density at the position assigned to dC25* (Fig. 6b). Density reappears for the PAM-distal segment from nucleotides 31–41, with nucleotides 33–41 forming dsDNA with the TS (Fig. 6b). The two resolved NTS segments are widely separated. The weak density at dC25* could reflect a mixture of cleavage products with different 3′ termini. Cleavage immediately after dC24* in a subset of molecules would be consistent with the putative nicking site identified in State III.

The cleaved PAM-proximal NTS product follows a continuous channel formed by RecA1, RecA2, and the CTD, with its newly generated 3′ end directed into the RecA1-RecA2 helicase channel (Fig. 6a). Comparison of the NTS conformations in State III and State IV revealed marked differences around the interface formed by Cas5, the Cas8 HB domain, and Cas2/3 (Fig. 6c). From the 5′ end to dT15*, the NTS follows a nearly identical path in both structures, with Cas8 HB and Cas5 sandwiching this segment in a similar manner (Figs. 6c and 6d). Beyond dT15*, however, the NTS trajectories diverge in opposite directions (Fig. 6d). In State III, nucleotides dG16*–dA19* form a continuous four-layer base stack that is further sandwiched by Trp296 of the Cas8 HB domain and Pro167 of Cas5, directing the downstream NTS toward the HD nuclease active site of Cas2/3 (Fig. 6e). In State IV, by contrast, the corresponding NTS segment bends sharply and is captured within the cleft between RecA2 and the CTD (Figs. 6d and 6e).

These structural differences suggest that, following NTS cleavage, the PAM-proximal NTS product dissociates from the Cas8 HB domain and is redirected toward the RecA2-CTD cleft before extending into RecA1. Superimposition of the State III and State IV structures further revealed a repositioning of Cas2/3 relative to the Csy complex by approximately 5 Å shift and 3° rotation, accompanying relocation of the cleaved NTS into the RecA1 and RecA2 domains (Fig. 6f).

Together, these observations indicate that NTS cleavage is followed by release of the PAM-proximal NTS product from the Cas8 HB domain and its redirection into the RecA1-RecA2 helicase channel.

## Discussion

Here, we provide a structural framework for the sequential events linking target DNA recognition to Cas2/3-mediated cleavage in the type I-F CRISPR-Cas system. Our structures spanning States I–IV support a stepwise model of CRISPR interference (Fig. 7). Target DNA binding induces Csy elongation and rotation of the Cas8 HB domain, creating a platform for Cas2/3 recruitment. Before recruitment, the NTS segment destined for initial cleavage remains flexible. Cas2/3 engagement stabilizes this segment as a bulge at the HD nuclease active site, with the Cas8 HB domain contributing to substrate positioning. Recruitment is also proposed to relieve RecA1 gate loop-mediated autoinhibition, allowing the NTS to access the HD active site. Following initial nicking, the PAM-proximal NTS product is released from its pre-cleavage contacts with the Cas8 HB domain and redirected into the channel formed by RecA1, RecA2, and the CTD. Based on previous studies of Cas3 and Cas2/3^23,41,42^, we propose that the helicase module subsequently reels in the NTS in an ATP-dependent manner, feeding it into the HD active site for processive 3′-to-5′ degradation. Together, these structures connect formation of the Cas2/3 recruitment platform with NTS bulge capture, initial nicking, and cleavage-product handover to the helicase.

**Figure 7.**
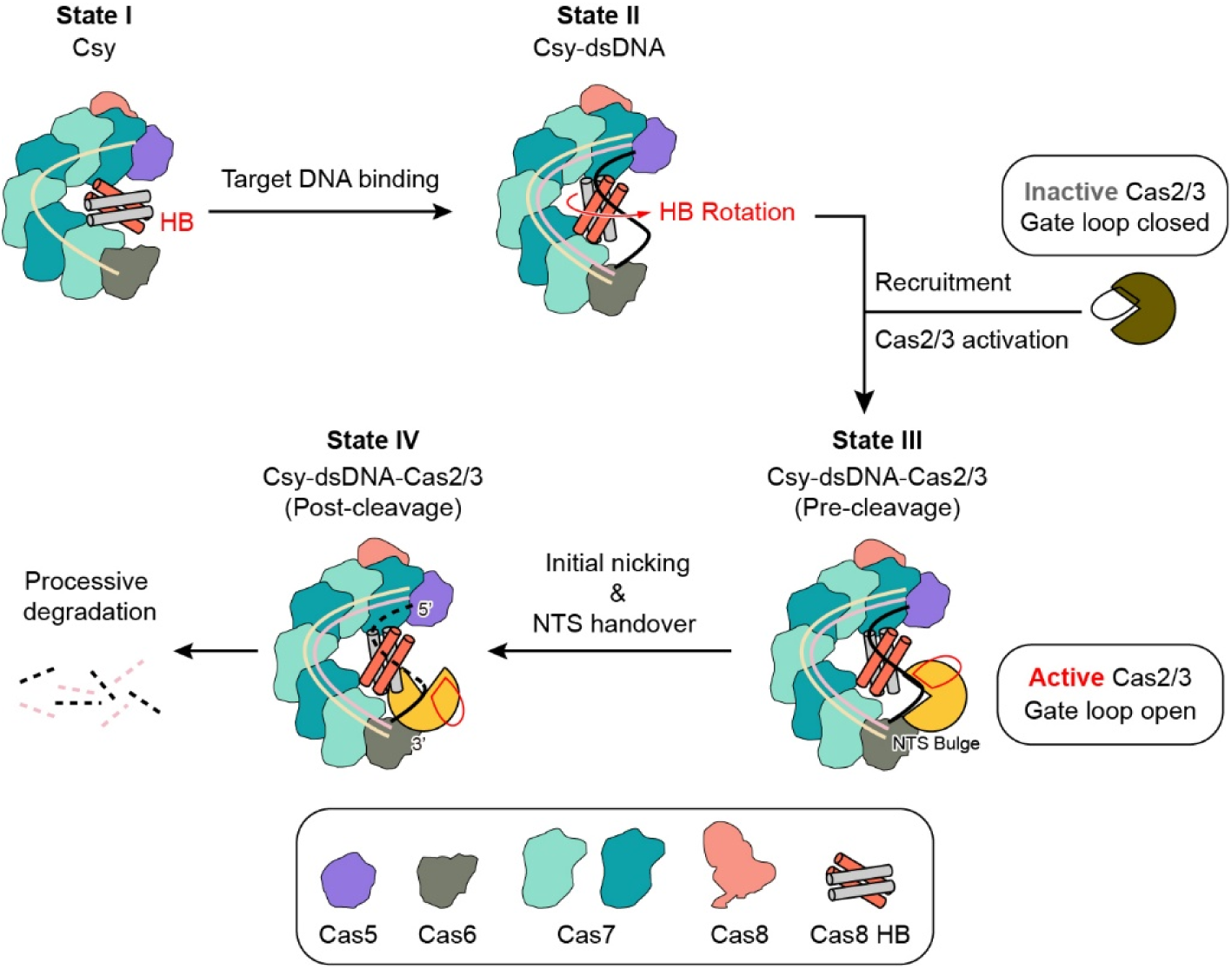
Proposed model of Cas2/3 recruitment, activation, and target DNA degradation in type I-F CRISPR-Cas interference. The model links target DNA recognition and Cas2/3 recruitment to NTS bulge capture, initial nicking, and cleavage-product handover to the helicase module. Recruitment-induced relief of gate loop-mediated autoinhibition and subsequent ATP-dependent processive degradation are proposed based on the present structures and previous studies. The crRNA, TS, and NTS are shown in wheat, pink, and black, respectively, and the RecA1 gate loop is shown in red.

Crystal structures of the type I-E Cas3-ssDNA complex revealed a closed helicase channel through which ssDNA extends toward the HD nuclease active site^23^. In these structures, ssDNA lies beneath the loop connecting the linker domain and the CTD. This arrangement raised a topological question of how the intact NTS within an R-loop could enter the closed channel without a free 3′ end. Subsequent structures established a helicase-bypassing route for initial NTS nicking in type I-E^15^ and resolved the NTS bulge captured at the HD active site in type I-C^26^. After nicking, density interpreted as the relocated NTS product was observed near the helicase entrance in type I-E, although its path within the channel remained unresolved^15^. A type I-A structure also captured the nicked PAM-proximal NTS segment threaded through the helicase toward the HD active site^27^. NTS engagement by the helicase was also observed in the Cas1-dependent type I-F system of ICP1, a phage infecting *V. cholerae*, where the NTS enters a channel enclosed by the CTD, linker, and RecA2 domains and extends across RecA1 toward the HD domain^33^. Our structures extend these findings by resolving both the NTS positioned for initial cleavage in the HD active site and the PAM-proximal cleavage product accommodated within the helicase channel of the VpCsy-DNA-Cas2/3 complex. The relatively weak density for the NTS bulge in State III raises the possibility that the bulge adopts slightly different conformations, allowing initial nicking at nearby phosphodiester bonds. This could account for the weak density at dC25* in State IV if the reconstruction contains cleavage products with different 3′ termini. In State IV, the product lies beneath the loop connecting the linker domain and the CTD (residues 1020–1029), consistent with loading of the newly generated 3′ end after initial nicking. Comparison of States III and IV reveals different NTS paths and accompanying changes in contacts with the Cas8 HB domain and Cas5. Together with the contacts stabilizing the product within the RecA1-RecA2-CTD channel, these observations provide a structural basis for how Cas2/3 initially nicks the NTS through a helicase-bypassing route and subsequently captures the cleavage product within its helicase channel.

The architecture of the Csy complex may help define the NTS region captured for initial cleavage. Type I-F Csy complexes adopt a pronounced curvature, with the TS following the curved Csy backbone and the NTS taking a distinct path on the opposite side of the Cas8 HB domain. By stabilizing the TS at the PAM-distal end of the R-loop, the Cas8 HB domain may also help prevent re-annealing of the TS and NTS. In the State II structure of VpCsy, NTS nucleotides 20–31 are disordered, whereas the flanking nucleotides dA19* and dA32* are stabilized by stacking interactions with Trp296 and Tyr324 of the Cas8 HB domain, respectively. These flanking nucleotides are separated by approximately 23 Å, suggesting that the 12 intervening nucleotides are longer than required to span this distance and can adopt flexible, non-extended conformations before Cas2/3 recruitment. These observations suggest that the Cas8 HB domain anchors the NTS on either side of the flexible region. Upon Cas2/3 recruitment, this region is captured and stabilized as a bulge within the HD nuclease domain, as observed in State III. Csy therefore appears to help define the region available for bulge formation, whereas interactions with Cas2/3 stabilize the bulge and position the scissile phosphodiester bond at the HD active site.

Cas8-Cas3 interactions play a central role in coupling target recognition to DNA cleavage in canonical type I CRISPR-Cas systems, although the interaction interfaces and regulatory mechanisms vary among subtypes. In type I-C, I-E, and I-F systems, full R-loop formation induces conformational changes in Cas8 that enable Cas3 recruitment^5,15,26^. In the type I-A system, Cas3 is constitutively associated with Cascade, but its HD active site remains occluded until full R-loop formation triggers allosteric activation^27^. Despite their mechanistic differences, these systems share a common regulatory principle: Cas3-mediated DNA degradation is licensed only after full R-loop formation, providing a safeguard against off-target DNA cleavage. In the structurally characterized type I-A, I-C, and I-E complexes, Cas3 binds near the PAM-proximal region, consistent with initial NTS nicking closer to the PAM than in VpCsy^15,26,27^. By contrast, Cas2/3 in VpCsy occupies a more PAM-distal position, where it contacts the Cas8 HB domain, which rotates upon target DNA binding. This arrangement places the HD active site near the flexible NTS region described above. Our structures show how this region forms a bulge that positions the phosphodiester bond between NTS nucleotides 24 and 25 for initial cleavage. Thus, the distinct Cas8-Cas2/3 arrangement provides a structural explanation for the more PAM-distal nicking region in VpCsy relative to those reported in type I-A, I-C, and I-E systems.

The PAM-distal placement of Cas2/3 in VpCsy is mediated by multiple contacts with Cas8 and Cas5. Previous studies proposed that the rotated Cas8 HB domain recruits Cas2/3 on the basis of its structural similarity to the anti-CRISPR protein AcrIF3 and mutational analysis of the predicted recruitment interface^5^. Our structures directly reveal the interaction between the Cas8 HB domain and the Cas2/3 CTD, supporting this model. These structures also reveal additional contacts between Cas5 and the Cas2/3 HD and linker domains, including engagement of the Cas2/3 anchor loop by both Cas5 and the Cas8 recruitment loop. These contacts may stabilize Cas2/3 in a position that allows the HD domain to capture the nearby NTS bulge. Although the interface residues are not identical between VpCas8 and PaeCas8, mutations at structurally corresponding positions in PaeCas8 have been shown to impair Cas2/3 recruitment. Together with our structural observations, these findings support a conserved role for the HB-mediated recruitment interface despite sequence divergence and reveal how additional Cas5 contacts contribute to the architecture of the Csy-Cas2/3 complex.

Comparison with noncanonical type I-F systems highlights the diversity of Cas2/3 recruitment mechanisms. In the ICP1 phage system, Cas8 lacks the HB domain, and a Cas1 dimer bridges the Csy complex and Cas2/3, with one Cas1 subunit contacting Cas6 and the other contacting Cas2/3^33^. In the type I-F2 system, which lacks Cas8 altogether, Cas7 subunits directly recruit Cas3^36^. These alternative recruitment interfaces contrast with the HB-mediated interface that we directly observed in our canonical type I-F structures. Despite these differences, target-induced elongation of the Csy complex provides a common means of controlling Cas2/3 recruitment in VpCsy and ICP1. In both systems, elongation relieves steric clashes that would otherwise prevent Cas2/3 binding. Thus, despite their distinct recruitment interfaces, both systems employ a similar strategy that couples target recognition to nuclease engagement. Our structures define this process in a canonical type I-F system and reveal how the recruited Cas2/3 captures the NTS for initial nicking and subsequently accommodates the cleavage product within its helicase channel.

## Acknowledgements

We thank Keishiro Uda and Osamu Kikko for their assistance with cryo-EM data collection and single-particle analysis. We also thank Hideko Inanaga for her technical assistance during the experiments. Cryo-EM data were collected by using EM01CT and EM02CT at SPring-8 (Hyogo, Japan), with the approval of the Japan Synchrotron Radiation Research Institute (JASRI) under proposal numbers 2021A2715, 2021B2548, 2022B2533, 2023B2531, and 2024B2523 to T.N. and 2023B0319 to K.I. This study was partially supported by the Research Support Project for Life Science and Drug Discovery (Basis for Supporting Innovative Drug Discovery and Life Science Research (BINDS)) from AMED under Grant Number JP25ama121001 (Support Number 4257).

## Author contributions

H.K. designed the research, performed biochemical and structural experiments, and wrote the manuscript; K.I. helped with research design and structural experiments; T.O. helped with cryo-EM data collection; C.G., C.C.G., and H.S. assisted with cryo-EM data collection. T.N. conceived the project, designed experiments, analyzed data, and wrote the manuscript. All authors contributed to writing and editing the manuscript.

## Methods

### Plasmid construction

For production of the *V. parahaemolyticus* 3259 Csy complex, the operon encoding Cas8, Cas5, Cas7, and Cas6 was synthesized by GeneArt (Thermo Fisher Scientific) and cloned into the multiple cloning site downstream of the SUMO-coding sequence in pE-SUMOproAmp using BsaI (New England Biolabs) and DNA Ligation Mix (TaKaRa) to yield pE-SUMOproAmp_VpCas8576. Internal BsaI recognition sites within the Csy operon were removed during DNA synthesis. A DNA fragment comprising the first two repeats and the first spacer of the *V. parahaemolyticus* 3259 CRISPR array was generated by annealing two partially complementary synthetic oligonucleotides, followed by DNA polymerase-mediated extension. The resulting fragment was cloned into the StuI and XhoI sites of a pCOLADuet-1 vector previously modified in our laboratory using NEBuilder HiFi DNA Assembly Master Mix (New England Biolabs) to yield pCOLADuet-1-StuI_VpcrRNA. Cloning at these sites removed the DNA sequences encoding the vector-derived ribosome-binding site and start codon, thereby excluding the corresponding sequences from the pre-crRNA transcript to minimize the potential for unintended secondary structure formation prior to processing.

For production of *V. parahaemolyticus* 3259 Cas2/3, the corresponding gene was synthesized by GeneArt (Thermo Fisher Scientific) and cloned into the EcoRI and XhoI sites of pCDFDuet-1 using NEBuilder HiFi DNA Assembly Master Mix to yield pCDFDuet-1_VpCas2/3.

For the *in vivo* plasmid interference assay, the operon encoding Cas8, Cas5, Cas7, and Cas6 and the CRISPR array were cloned into the NdeI-XhoI and StuI-NotI sites, respectively, of the modified pCOLADuet-1 vector using NEBuilder HiFi DNA Assembly Master Mix to yield pCsy. DNA fragments containing a 5′-CC-3′ PAM and a protospacer were cloned into the BamHI and EcoRI sites of pUC18 using NEBuilder HiFi DNA Assembly Master Mix to yield pTarget.

Site-directed mutagenesis was performed by inverse PCR, followed by DpnI digestion to remove the template plasmid. The PCR products were phosphorylated using T4 polynucleotide kinase (New England Biolabs) and circularized by ligation using Mighty Mix (TaKaRa). The sequences of the resulting plasmids were verified by Sanger sequencing.

### Preparation of VpCsy complex

*E. coli* BL21-Gold(DE3)-CodonPlus-RIL competent cells were co-transformed with pE-SUMOproAmp_VpCas8576 and pCOLADuet-1-StuI_VpcrRNA. The cells were cultured at 37°C in LB medium supplemented with 50 μg/mL ampicillin, 50 μg/mL kanamycin, and 34 μg/mL chloramphenicol until the OD₆₀₀ reached 0.5. Protein expression was then induced by adding isopropyl β-D-1-thiogalactopyranoside (IPTG) to a final concentration of 0.1 mM. The culture was incubated overnight at 20°C, and the cells were harvested by centrifugation at 5480 × *g* for 10 min, followed by 8000 × *g* for 10 min. The cell pellet was resuspended in lysis buffer containing 20 mM Tris-HCl, pH 8.0, 300 mM NaCl, 7 mM 2-mercaptoethanol, 1 mM benzamidine, and 1 mM phenylmethylsulfonyl fluoride (PMSF). The cells were lysed by sonication on ice at 30% amplitude with 1 sec on/1 sec off pulses for a total on-time of 4 min. The lysate was clarified by sequential centrifugation at 7,600 × *g* for 15 min and 14,300 × *g* for 15 min. The resulting supernatant was applied to Ni-NTA agarose (Qiagen), and bound proteins were eluted with buffer containing 20 mM Tris-HCl, pH 8.0, 300 mM NaCl, 250 mM imidazole, and 7 mM 2-mercaptoethanol. The His-SUMO tag was cleaved with Ulp1 protease, and the sample was reapplied to Ni-NTA agarose to remove the cleaved tag. The flow-through fraction was collected and loaded onto a 5 mL HiTrap Heparin HP column (Cytiva). The Csy complex was eluted with a linear gradient of 100–1,000 mM NaCl in buffer containing 20 mM Tris-HCl, pH 8.0, and 7 mM 2-mercaptoethanol. The complex was further purified by size-exclusion chromatography (SEC) on a Superdex 200 Increase 10/300 GL column (Cytiva) equilibrated with buffer containing 20 mM Tris-HCl, pH 8.0, 100 mM NaCl, and either 7 mM 2-mercaptoethanol for *in vitro* assays or 1 mM dithiothreitol (DTT) for cryo-EM data collection. Mutant Csy complexes were purified using the same procedure as that used for the wild-type complex.

### Preparation of VpCas2/3

*E. coli* BL21-Gold(DE3)-CodonPlus-RIL competent cells were transformed with pCDFDuet-1_VpCas2/3. The cells were cultured at 37°C in LB medium supplemented with 50 μg/mL streptomycin and 34 μg/mL chloramphenicol. When the OD₆₀₀ reached 0.5, protein expression was induced by adding IPTG to a final concentration of 0.1 mM, and the culture was incubated at 20°C for approximately 18 h. The cells were harvested by centrifugation at 5480 × *g* for 10 min, followed by 8000 × *g* for 10 min. The cell pellet was resuspended in the lysis buffer and lysed by sonication under the same conditions as those used for the Csy complex. The lysate was clarified by sequential centrifugation at 7,600 × *g* for 15 min and 14,300 × *g* for 15 min. The resulting supernatant was applied to Ni-NTA agarose (Qiagen), and bound proteins were eluted as described above. The eluate was loaded onto a 1 mL HiTrap Heparin HP column (Cytiva), and Cas2/3 was eluted with a linear gradient of 300–1,000 mM NaCl in buffer containing 20 mM Tris-HCl, pH 8.0, and 7 mM 2-mercaptoethanol. Cas2/3 was further purified by SEC on a Superdex 200 Increase 10/300 GL column (Cytiva) equilibrated with buffer containing 20 mM Tris-HCl, pH 8.0, 300 mM NaCl, and either 7 mM 2-mercaptoethanol for *in vitro* assays or 1 mM DTT for cryo-EM data collection.

### Reconstitution of VpCsy-dsDNA and VpCsy-dsDNA-Cas2/3 complexes

To prepare target dsDNA containing a 5′-CC-3′ PAM, complementary 75-mer oligonucleotides were annealed in hybridization buffer containing 20 mM Tris-HCl, pH 8.0, 100 mM NaCl, and 5 mM MgCl₂ by heating at 95°C for 5 min, followed by gradual cooling to room temperature over 1 h. The resulting dsDNA and the Csy complex were mixed at a molar ratio of 1.5:1, and the mixture was incubated at 37°C for 5 min to reconstitute the complex. The reconstituted Csy-dsDNA complex was purified by SEC on a Superdex 200 Increase 10/300 GL column (Cytiva) equilibrated with buffer containing 20 mM Tris-HCl, pH 8.0, 100 mM NaCl, and 1 mM DTT.

To reconstitute the Csy-dsDNA-Cas2/3 complex, the purified Csy-dsDNA complex and Cas2/3 were mixed at a molar ratio of 1:3 and incubated at 37°C for 15 min in buffer containing 20 mM Tris-HCl, pH 8.0, 300 mM NaCl, 7 mM 2-mercaptoethanol, and 5% glycerol. VpCas2/3 was used for complex reconstitution on the day of purification. The reconstituted complex was purified by SEC on a Superdex 200 Increase 10/300 GL column (Cytiva) equilibrated with buffer containing 20 mM Tris-HCl, pH 8.0, 300 mM NaCl, and 1 mM DTT.

### Cryo-EM sample preparation and data collection

Cryo-EM grids were prepared using different sample concentrations, detergents, and glow-discharge and blotting conditions suitable for each complex. For all samples, 3 μL was applied to glow-discharged Quantifoil R1.2/1.3 200-mesh copper grids.

For the DNA-unbound VpCsy complex (State I), the sample was prepared at 10 mg/mL with 8 mM CHAPS, and the grids were glow-discharged at 15 Pa and 10 mA for 20 s. For the target DNA-bound VpCsy-dsDNA complex (State II), the sample was prepared at 15 mg/mL with 8 mM CHAPS, and the same glow-discharge conditions were used. For the VpCsy-dsDNA-Cas2/3 complex (States III and IV), the sample was prepared at 12 mg/mL with 0.01% (v/v) NP-40, and the grids were glow-discharged at 20 Pa and 20 mA for 45 s.

All grids were blotted at 8°C and 100% humidity with a blot force of 0 using a Vitrobot Mark IV (Thermo Fisher Scientific). The blotting time was 2 s for State I and States III and IV, and 3 s for State II. The grids were then plunge-frozen in liquid ethane cooled with liquid nitrogen.

Cryo-EM data were collected at SPring-8 using a CRYO ARM 300^TM^ (JEOL) equipped with an in-column Omega energy filter and a K3 direct electron detector (Gatan) operated in counting mode. Hole detection was performed using YoneoLocr^43^, and automated data acquisition was carried out using SerialEM^44^ at a nominal magnification of 60,000×, corresponding to a calibrated pixel size of 0.752 Å. Movies were recorded as 50-frame stacks with a total electron exposure of 50 e⁻/Å², using a target defocus range of −1.4 to −1.6 μm, and a 5 × 5 beam-image shift pattern.

### Image processing

The single-particle analysis workflows for the DNA-unbound VpCsy complex (State I), the VpCsy-dsDNA complex (State II), and the VpCsy-dsDNA-Cas2/3 complex (States III and IV) are shown in Supplementary Figures S1, S2, and S5, respectively.

For State I, a total of 6,800 movies were motion-corrected and subjected to contrast transfer function (CTF) estimation in CryoSPARC Live v4.6.1^45^, and micrographs with CTF fit resolutions worse than 4 Å were excluded in CryoSPARC. Initial particle picking from 50 micrographs was performed using Blob Picker (diameter 150–200 Å). Particles containing contaminants were excluded, and the remaining particles were extracted with a box size of 320 pixels and Fourier-cropped to 80 pixels. Using the resulting 2D class averages as templates, particle picking was then conducted across all micrographs with CTF fit resolutions better than 4 Å, using a particle diameter of 180 Å. The extracted particles were subjected to 2D classification. Initial 3D models were generated by Ab-Initio Reconstruction in CryoSPARC with K=3, using an initial resolution of 35 Å and a maximum resolution of 12 Å. Using these references, contaminating particles were removed through Heterogeneous Refinement with a box size of 128 pixels. The major class was re-extracted at a box size of 360 pixels and refined by Non-Uniform Refinement. Particles were then optimized by CTF Refinement and subjected to Reference-Based Motion Correction. The particles were further classified by 3D Classification without alignment (K=2) to exclude a class corresponding to Csy complex artifacts lacking Cas6. The class in which the Cas8 HB domain and Cas6 were most clearly resolved was refined by Non-Uniform Refinement. The regions comprising the Cas8 HB domain, Cas5, Cas7.1, and Cas6 were further refined by Local Refinement.

For State II, a total of 7,411 movies were motion-corrected and subjected to contrast transfer function (CTF) estimation in CryoSPARC Live v4.6.1^45^. Streaming 2D Classification was performed and particles containing contaminants were excluded. Using the resulting 2D class averages as templates, initial 3D models were generated by Ab-Initio Reconstruction in CryoSPARC with K=3, using an initial resolution of 35 Å and a maximum resolution of 12 Å. Using these references, contaminating particles were removed through Heterogeneous Refinement with a box size of 128 pixels. The major class was re-extracted at a box size of 480 pixels and refined by Homo Refinement. Particles were then optimized by CTF Refinement and subjected to Reference-Based Motion Correction. The particles were further classified by 3D Classification without alignment (K=2) to exclude a class corresponding to Csy complex artifacts lacking Cas6. The class in which the Cas8 HB domain and Cas6 were most clearly resolved was refined by Non-Uniform Refinement. The regions comprising the Cas8 HB domain and Cas6 were further refined by Local Refinement.

For States III and IV, a total of 8,151 movies were imported into RELION-4.0.1^46^ for motion correction. Motion-corrected micrographs were transferred to CryoSPARC v4.6.1^45^ for contrast transfer function (CTF) estimation, and micrographs with CTF fit resolutions worse than 5 Å were excluded. Initial particle picking from 50 micrographs was performed using Blob Picker (diameter 150–200 Å). Particles with contaminants were excluded, and the remaining particles were extracted with a box size of 400 pixels and Fourier-cropped to 80 pixels. Using the 2D class averages as templates, particle picking was conducted across all micrographs with a particle diameter of 230 Å. Subsequently, the extracted particles were subjected to 2D classification. Initial 3D models were generated by Ab-Initio Reconstruction in CryoSPARC with K=3, using an initial resolution of 35 Å and a maximum resolution of 12 Å. Using these references, contamination was removed through heterogeneous refinement with a box size of 128 pixels. The major class was refined using Non-Uniform Refinement, after particles were re-extracted at 350 pixels. Particles optimized by CTF refinement were then imported into RELION-4.0.1 for Bayesian polishing. Particles were re-imported into CryoSPARC and subjected to 3D classification without alignment (K=5) to exclude classes corresponding to the Csy complex alone. The resulting particles were subjected to 3D Variability to resolve conformational heterogeneity within the Cas2/3 module. Particles were then separated to the dataset into two populations reflecting distinct classes of nucleic acid engagement by Cas2/3: one in which the substrate was engaged by the HD nuclease domain (State III), and another in which it was engaged by the RecA domain (State IV).

For State III, particles were re-imported into RELION-4.0.1 and subjected to focused 3D classification without alignment (T=32) using a mask encompassing the Cas2/3 region, and the class with the strongest density was selected. Particle signal was then reverted to that of the intact complex. After re-importing into CryoSPARC, particles were subjected to focused 3D classification using a small mask encompassing the HD domain to resolve nucleic acid engagement, and the class showing nucleic acid density within the HD domain was selected. To exclude particles in which the nucleic acid was additionally engaged by the RecA domain, the particles were further classified using a smaller mask encompassing the Cas2/3 RecA region, and the class lacking RecA-associated nucleic acid density was selected and refined by Non-Uniform Refinement. For State IV, particles were subjected to focused 3D classification using a small mask encompassing the Cas2/3 RecA region, and the class exhibiting nucleic acid density within the RecA domain was selected and refined by Non-Uniform Refinement. For both States III and IV, the regions comprising Cas2/3 and Cas6 were further refined by Local Refinement.

For all states, the resolution of each map was estimated using the gold-standard Fourier shell correlation (FSC) criterion^47^ of 0.143, and local resolution was estimated using the Local Resolution Estimation tool.

### Model building and refinement

Atomic models of each complex were built using structural models generated by AlphaFold3^48^. The predicted model of each Cas protein subunit was first fitted into the cryo-EM maps using the “Fit in Map” function in UCSF ChimeraX^49^ and then manually adjusted in Coot^50^. Nucleic acid models were built manually in Coot^50^. The models were iteratively refined by real-space refinement in Phenix^51^ and manual rebuilding in Coot^50^. Model quality was assessed using MolProbity^52^ implemented in Phenix^53^. The final refinement and validation statistics are summarized in Table S1. All structural figures were prepared using UCSF ChimeraX^49^ and PyMOL (https://www.pymol.org/).

### *In vivo* plasmid interference assay

Three plasmids (pCsy, pCDFDuet-1_VpCas2/3, and pTarget) were co-transformed into *E. coli* BL21-Gold(DE3) competent cells, and the cells were plated on LB agar plates containing kanamycin (Kan), streptomycin (St), and ampicillin (Amp). After overnight incubation at 37°C, a single colony was picked and inoculated into LB medium supplemented with three antibiotics. Cultures were grown until the OD₆₀₀ reached 0.5, 3 mL of the culture was harvested, and the cells were resuspended in 2 mL LB containing kanamycin and streptomycin, and IPTG were added at a final concentration of 0.1 mM. The cultures were then incubated at 37°C for 1 h to induce expression. After induction, the cultures were serially diluted, and 100 μL of each dilution was plated on LB agar plates containing either Kan-St-Amp or Kan-St. After incubation at 37°C for more than 15 h, colonies on each plate were counted. The depletion ratio was calculated by dividing the number of colonies on Kan-St plates by the number of colonies on Kan-St-Amp plates.

### *In vitro* DNA cleavage assay

dsDNA substrate was prepared by annealing the TS and a 5′-Cy5-labeled NTS at a molar ratio of 1:1. DNA cleavage reactions were performed by incubating 200 nM Csy complex, 400 nM Cas2/3, and 20 nM Cy5-labeled 75-bp dsDNA substrate at 37°C in reaction buffer containing 20 mM Tris-HCl, pH 8.0, 200 mM NaCl, 5% glycerol, 7 mM 2-mercaptoethanol, 5 mM MgCl₂, 75 μM NiSO₄, 5 mM CaCl₂, and 1 mM ATP. Reactions were carried out in a total volume of 5 μL for 0, 1, 3, 5, 10, 30, or 60 min and quenched by adding 5 μL of stop solution containing 99% formamide and heating at 98°C for 15 min. Cleavage products were separated by electrophoresis on 15% polyacrylamide gels containing 8 M urea and visualized using a Typhoon phosphorimager (GE Healthcare).

### Electrophoretic mobility shift assay

Cy5-labeled 75-bp dsDNA was prepared as described above. Binding reactions containing 20 nM dsDNA and increasing concentrations of the Csy complex (0, 31.3, 62.5, 125, 250, 500, or 1,000 nM) were incubated at 37°C for 15 min in reaction buffer containing 20 mM Tris-HCl, pH 8.0, 200 mM NaCl, and 7 mM 2-mercaptoethanol. The reaction mixtures were subjected to electrophoresis on 6% native polyacrylamide gels, and Cy5-labeled DNA was visualized using a Typhoon phosphorimager (GE Healthcare).

To assess Cas2/3 binding to the Csy-dsDNA complex, 20 nM Cy5-labeled dsDNA was pre-incubated with 200 nM Csy complex at 37°C for 15 min in reaction buffer containing 20 mM Tris-HCl, pH 8.0, 200 mM NaCl, 5% glycerol, 7 mM 2-mercaptoethanol, 5 mM MgCl₂, 75 μM NiSO₄, 5 mM CaCl₂, and 1 mM ATP. Cas2/3 was then added to final concentrations of 0, 100, 200, 400, or 800 nM, and the mixtures were incubated at 37°C for 10 min. The reaction mixtures were subjected to electrophoresis on 4.5% native polyacrylamide gels, and Cy5-labeled DNA was visualized as described above.

