## Supplemental information for "Structural basis for Cas2/3 recruitment and nuclease-to-helicase DNA handover in type I-F CRISPR-Cas"

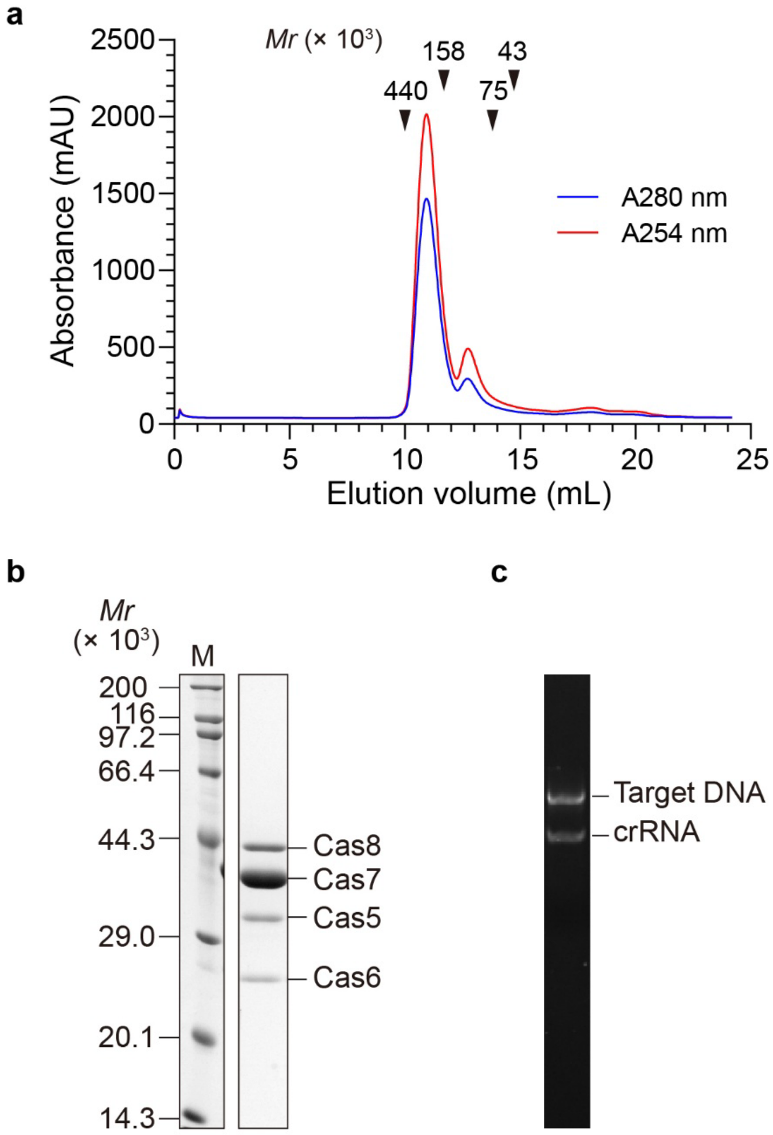


**Supplementary Figure S1.** Preparation of the target DNA-bound VpCsy complex.
**(a)** Size-exclusion chromatography profile of the target DNA-bound VpCsy complex obtained using a Superdex 200 Increase 10/300 GL column. Arrowheads indicate the elution positions of molecular mass standards: ferritin (440 kDa), aldolase (158 kDa), conalbumin (75 kDa), and ovalbumin (43 kDa). Absorbance at 254 and 280 nm is shown in red and blue, respectively. **(b)** SDS-PAGE analysis of the purified complex, showing the presence of Cas8, Cas7, Cas5, and Cas6. M, molecular mass markers. **(c)** Denaturing PAGE analysis of the purified complex, showing the presence of crRNA and target DNA.

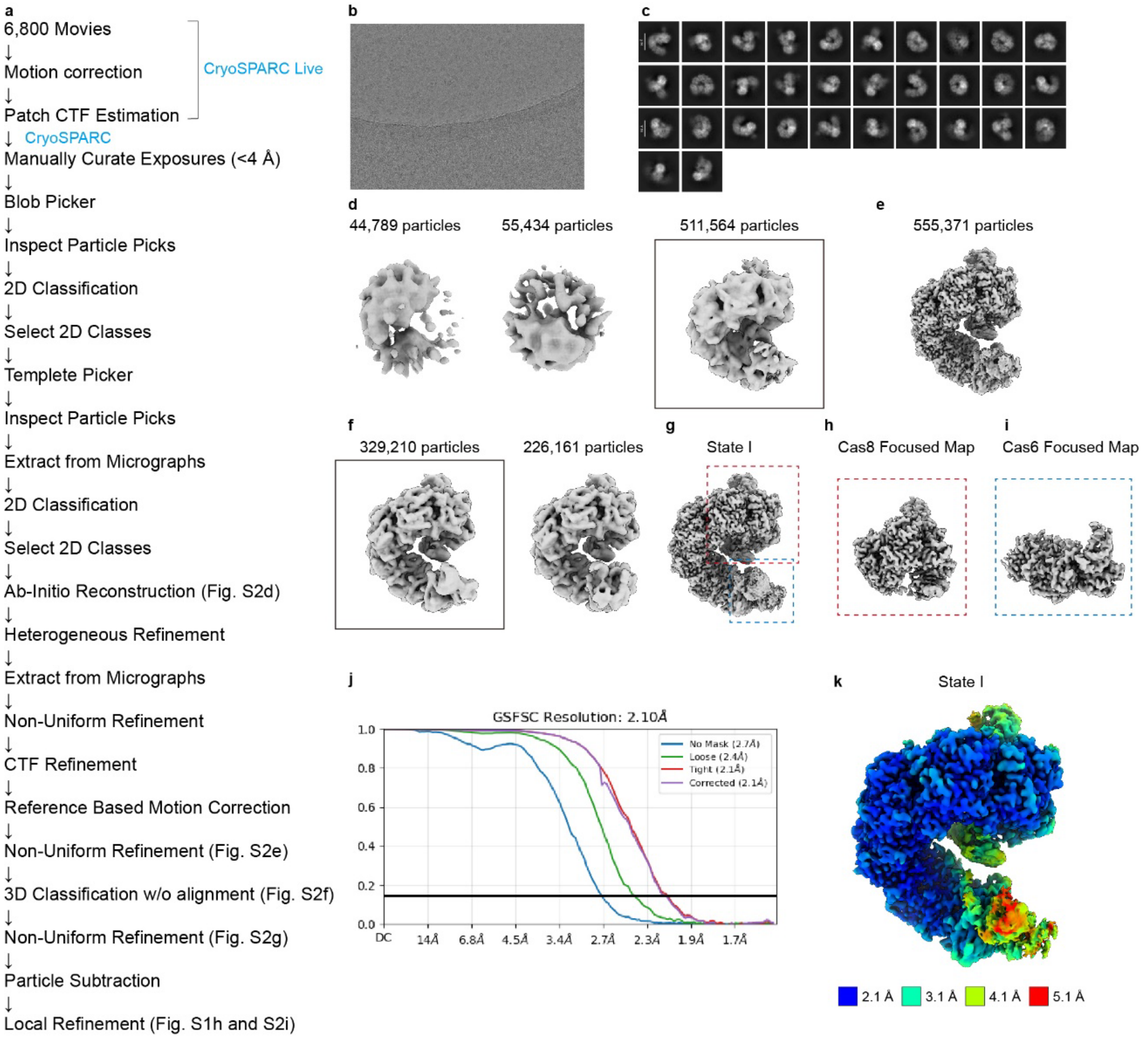


**Supplementary Figure S2.** Single-particle analysis of VpCsy complex alone (State I). **(a)** Workflow of single-particle analysis. **(b)** Representative motion-corrected micrograph. **(c)** Selected 2D class-averages for Ab-Initio Reconstruction. **(d)** Maps from Ab-Initio Reconstruction. The map enclosed in the box denotes the class selected for subsequent analysis. **(e)** 3D maps classified by multi-reference refinement through heterogeneous refinement. **(f)** Maps classified by 3D classification without alignment in CryoSPARC. The 3D map enclosed in the box denotes the class selected for further analysis. **(g)** The 3D Map refined by Non-Uniform Refinement in CryoSPARC. Dashed boxes indicate the region of the focused mask generated for the subsequent job. **(h, i)** Focused maps around Cas8 and Cas6, respectively. **(j)** Gold-standard Fourier shell correlation (FSC) curve. The solid horizontal line indicates the FSC at 0.143. **(k)** Estimated local resolution colored on the map.

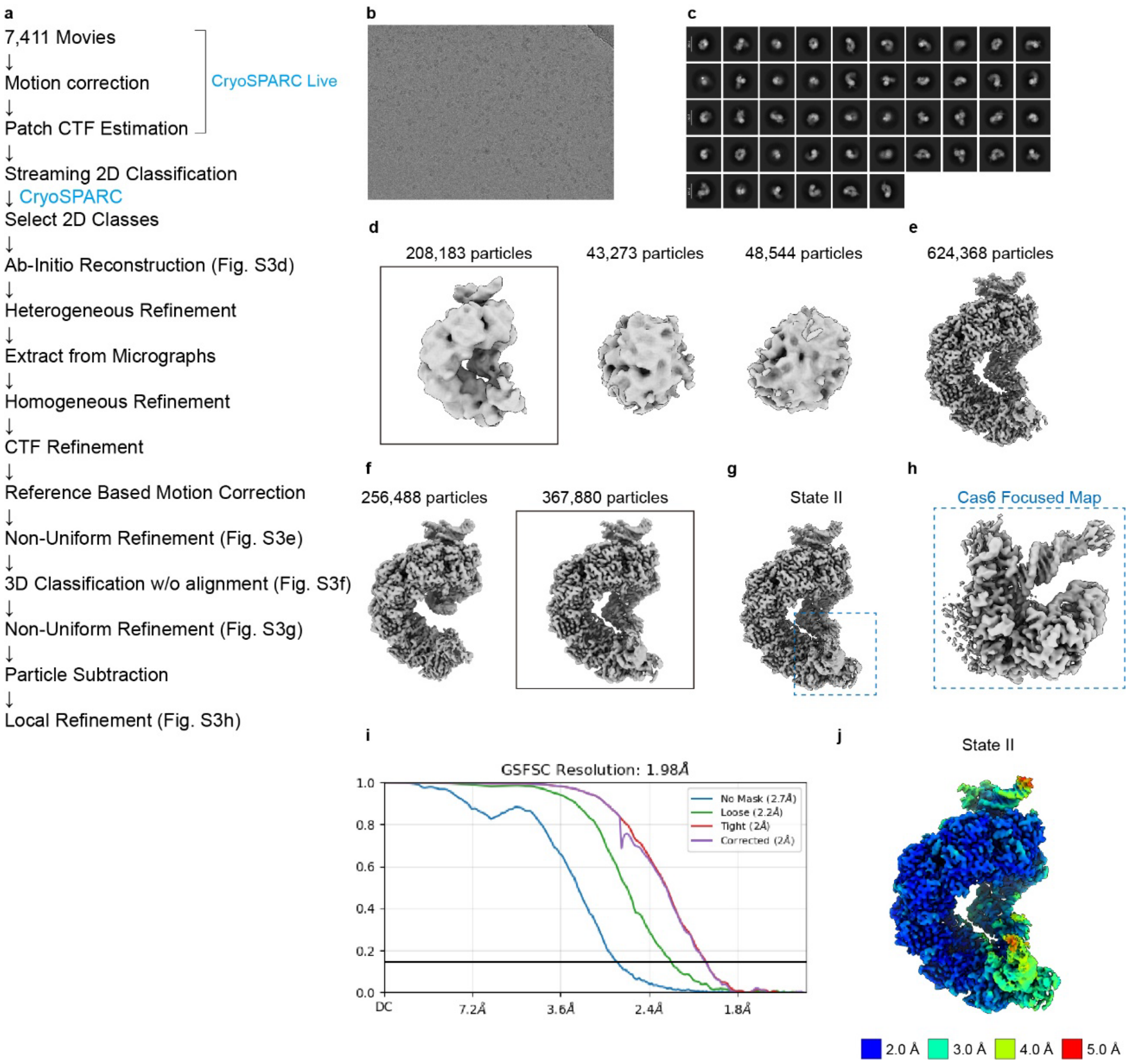


**Supplementary Figure S3.** Single-particle analysis of target DNA-bound VpCsy complex (State II).

**(a)** Workflow of single-particle analysis. **(b)** Representative motion-corrected micrograph. **(c)** Selected 2D class-averages for Ab-Initio Reconstruction. **(d)** Maps from Ab-Initio Reconstruction. The map enclosed in the box denotes the class selected for subsequent analysis. **(e)** 3D maps classified by multi-reference refinement through heterogeneous refinement. **(f)** Maps classified by 3D classification without alignment in CryoSPARC. The 3D map enclosed in the box denotes the class selected for further analysis. **(g)** The 3D Map refined by Non-Uniform Refinement in CryoSPARC. Dashed box indicates the region of the focused mask generated for the subsequent job. **(h)** Focused map around Cas6. **(i)** Gold-standard Fourier shell correlation (FSC) curve. The solid horizontal line indicates the FSC at 0.143. **(j)** Estimated local resolution colored on the map.

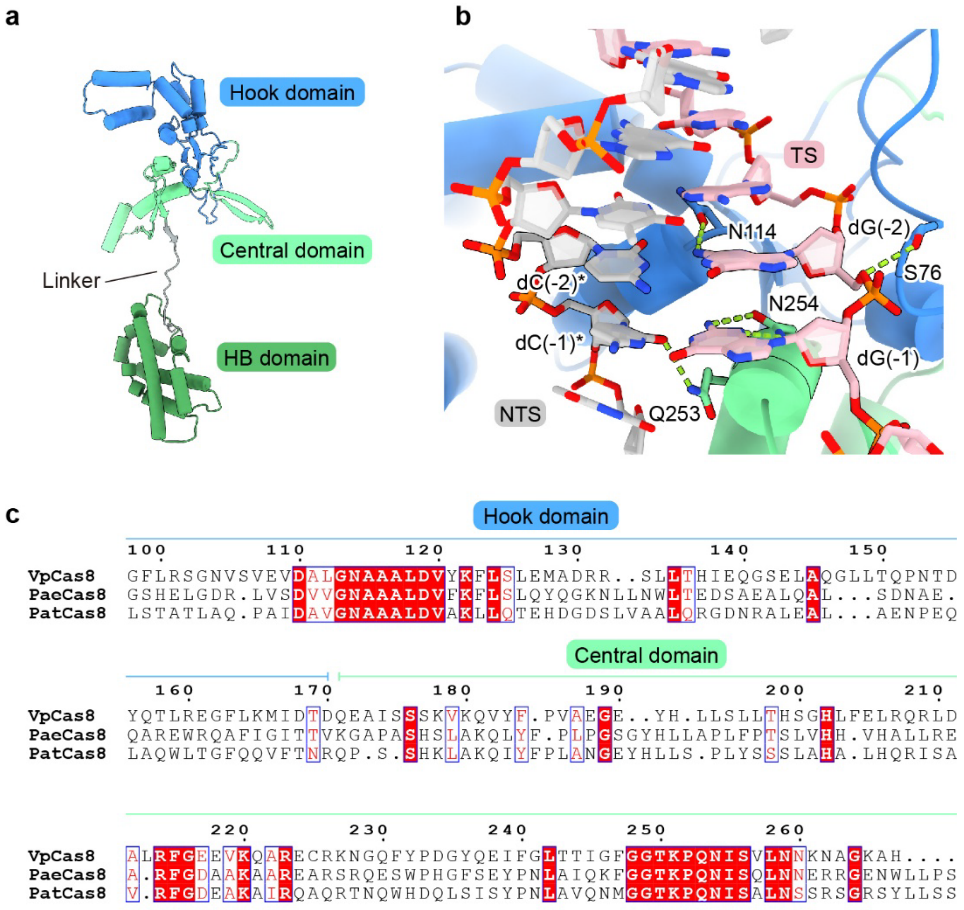


**Supplementary Figure S4.** PAM recognition by Cas8 within the VpCsy complex. **(a)** Overall structure of VpCas8 in State II. The hook, central, and helical bundle (HB) domains are colored marine blue, light green, and forest green, respectively. The linker segment is colored gray. **(b)** Close-up view of PAM recognition by VpCas8 in State II. The target DNA and selected amino acid residues involved in PAM recognition are shown as sticks. Nucleotides at PAM positions -2 and -1 are labeled. The TS and NTS are colored pink and gray, respectively. Hydrogen bonds are indicated by green dashed lines. VpCas8 domains are colored as in **a**. **(c)** Amino acid sequence alignment of the PAM recognition regions of VpCas8 and the corresponding regions of PaeCas8 and PatCas8. VpCas8 residue numbers are indicated above the alignment. The hook and central domain regions are indicated by horizontal lines colored as in **a**.

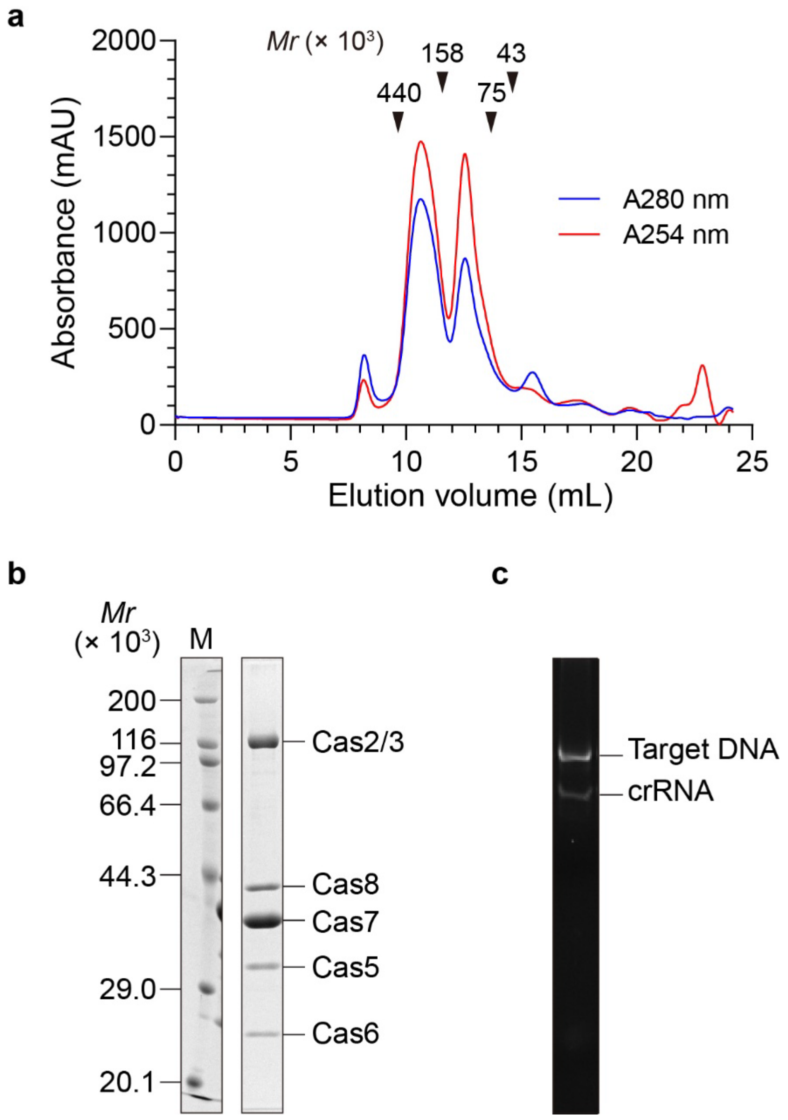


**Supplementary Figure S5.** Preparation of the VpCsy-DNA-Cas2/3 complex.

**(a)** Size-exclusion chromatography profile of the VpCsy-DNA-Cas2/3 complex obtained using a Superdex 200 Increase 10/300 GL column. Arrowheads indicate the elution positions of molecular mass standards: ferritin (440 kDa), aldolase (158 kDa), conalbumin (75 kDa), and ovalbumin (44 kDa). Absorbance at 254 and 280 nm is shown in red and blue, respectively. **(b)** SDS-PAGE analysis of the purified complex, showing the presence of Cas2/3, Cas8, Cas7, Cas5, and Cas6. M, molecular mass markers. **(c)** Denaturing PAGE analysis of the purified complex, showing the presence of crRNA and target DNA.

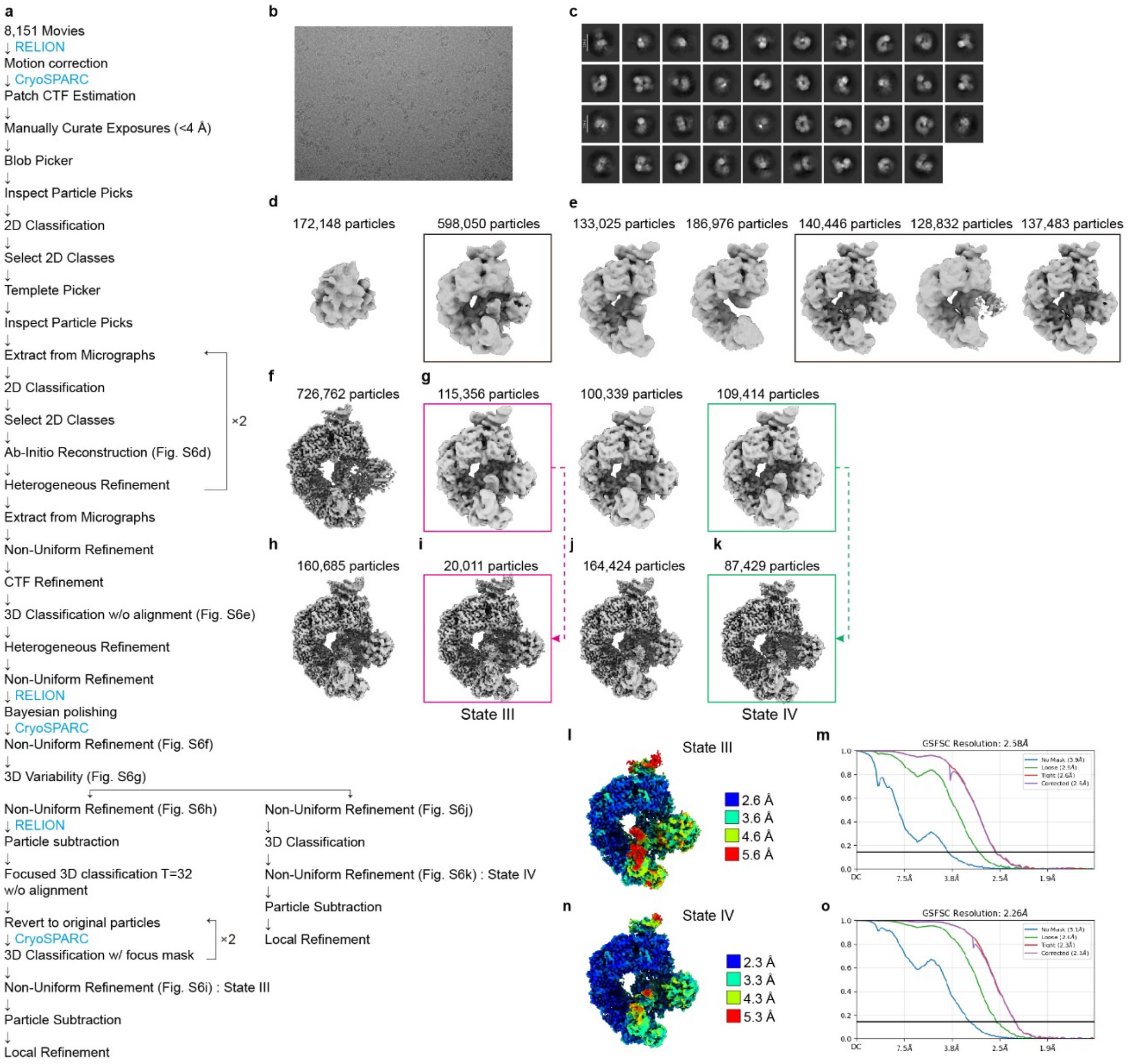


**Supplementary Figure S6.** Single-particle analysis of the VpCsy-DNA-Cas2/3 complexes (State III and State IV).

**(a)** Workflow of single-particle analysis. **(b)** Representative motion-corrected micrograph. **(c)** Selected 2D class-averages for Ab-Initio Reconstruction. **(d)** Maps from Ab-Initio Reconstruction. The map enclosed in the box denotes the class selected for subsequent analysis. **(e)** Maps classified by 3D classification without alignment in CryoSPARC. **(f)** The 3D Map refined by Non-Uniform Refinement in CryoSPARC. **(g)** 3D classes obtained by 3D variability analysis in CryoSPARC. Colored boxes indicate the individual states. **(h, i)** The 3D Maps refined by Non-Uniform Refinement in CryoSPARC (State III). **(j, k)** The 3D Maps refined by Non-Uniform Refinement in CryoSPARC (State IV). **(l, n)** Estimated local resolution colored on the map. **(m, o)** Gold-standard Fourier shell correlation (FSC) curve. The solid horizontal line indicates the FSC at 0.143.

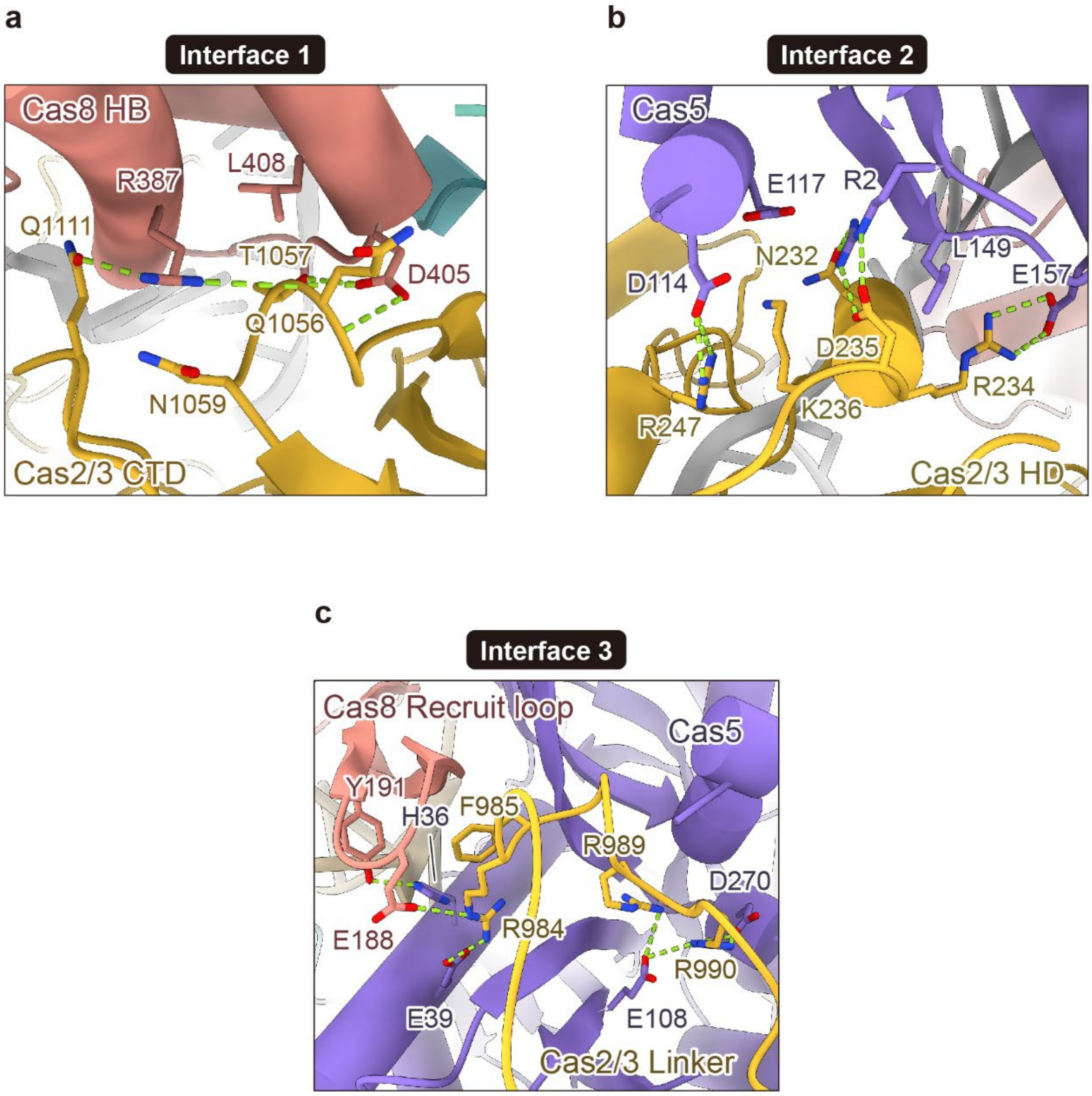


**Supplementary Figure S7.** Interactions between VpCsy and VpCas2/3 in State III.
**(a)** Close-up view of interface 1 between the Cas8 HB domain and the Cas2/3 CTD. **(b)** Close-up view of interface 2 between Cas5 and the Cas2/3 HD domain. **(c)** Close-up view of interface 3 involving Cas5, the nuclease recruitment loop of the Cas8 central domain, and the anchor loop of the Cas2/3 linker domain. In **a–c**, Cas2/3, Cas5, and Cas8 are colored yellow, purple, and salmon, respectively. Residues involved in the interactions are shown as sticks. Hydrogen bonds are indicated by green dashed lines.

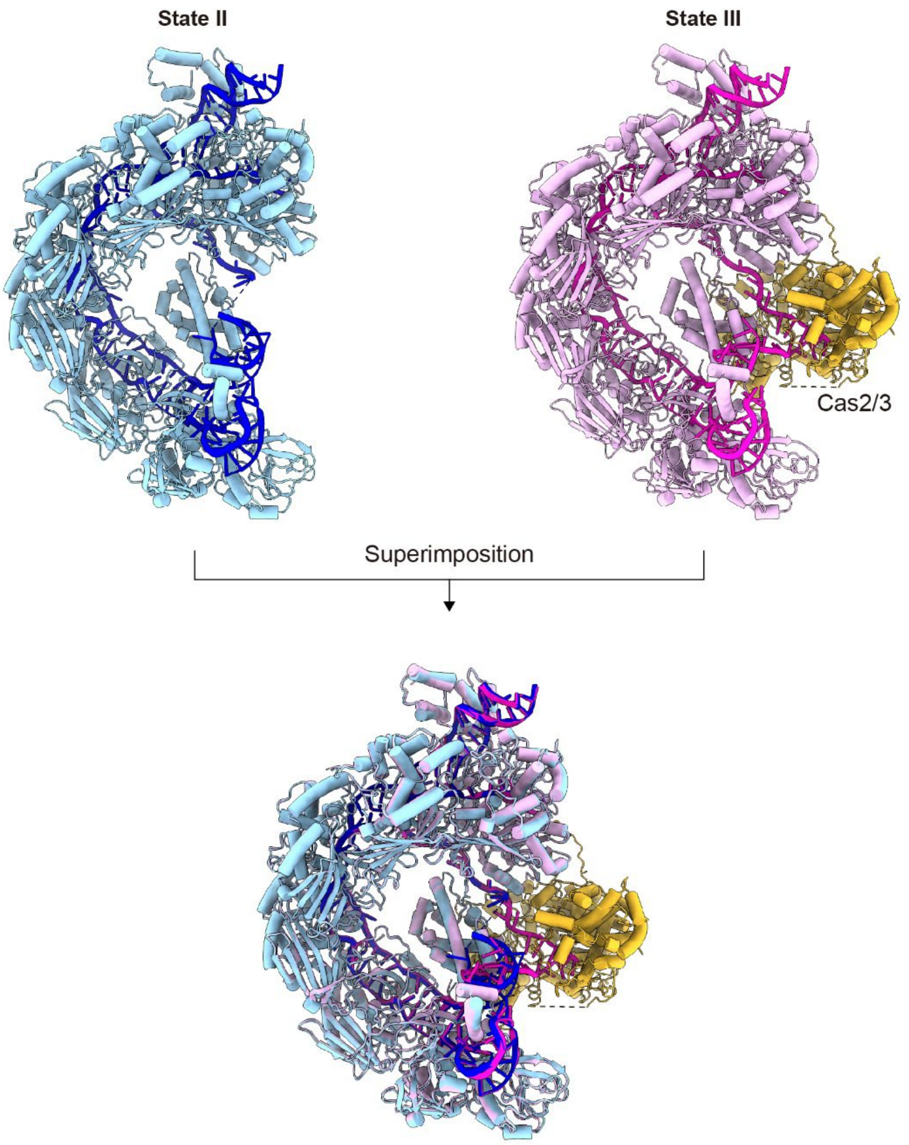


**Supplementary Figure S8.** Preorganized Cas2/3 binding site in the target DNA-bound Csy complex.

Overall structures of the target DNA-bound VpCsy complex in State II (upper left) and the VpCsy-DNA-Cas2/3 complex in State III (upper right), with their superimposition shown below. Protein subunits of the Csy complex are colored light blue in State II and pink in State III. Nucleic acids are colored blue in State II and magenta in State III. Cas2/3 is colored yellow. All structures are shown as cartoons.

a

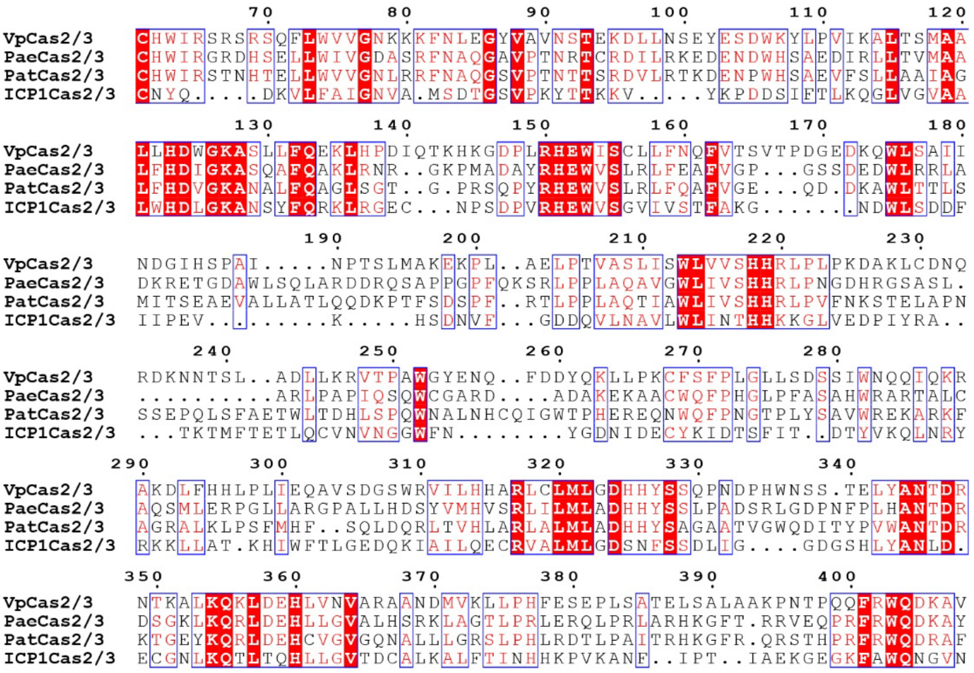


b

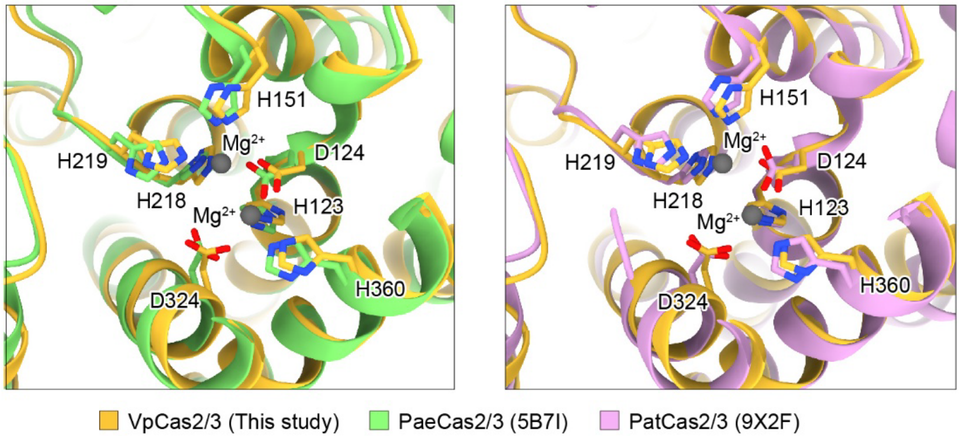


**Supplementary Figure S9.** Conservation of the Cas2/3 HD active site.

**(a)** Amino acid sequence alignment of HD domain from VpCas2/3, PaeCas2/3, PatCas2/3, and ICP1 Cas2/3. VpCas2/3 residue numbers are indicated above the alignment. **(b)** Structural comparison of the HD active sites. VpCas2/3 in State III is superimposed with PaeCas2/3 (PDB ID: 5B7I; left) and PatCas2/3 (PDB ID: 9X2F; right). VpCas2/3, PaeCas2/3, and PatCas2/3 are colored yellow, light green, and salmon, respectively. Conserved histidine and aspartate residues forming the active site are shown as sticks. Mg2+ ions observed in VpCas2/3 structure are shown in dark gray spheres. Residue numbers are those of VpCas2/3.

**Table 1. Cryo-EM data collection, refinement, and validation statistics**

| | State I | State II | State III | State IV |
|---|---|---|---|---|
| **Data collection and processing** | | | | |
| Microscope | CRYOARM™ 300 | CRYOARM™ 300 | CRYOARM™ 300 | CRYOARM™ 300 |
| Detector | Gatan K3 camera | Gatan K3 camera | Gatan K3 camera | Gatan K3 camera |
| Automation software | SerialEM | SerialEM | SerialEM | SerialEM |
| Nominal Magnification | 60,000 | 60,000 | 60,000 | 60,000 |
| Acceleration Voltage (kV) | 300 | 300 | 300 | 300 |
| Total electron exposure ($e^-$/Å$^2$) | 50.25 | 50.03 | 49.84 | 49.84 |
| Defocus range (μm) | −1.4 to −1.6 | −1.4 to −1.6 | −1.4 to −1.6 | −1.4 to −1.6 |
| Pixel size (Å) | 0.752 | 0.752 | 0.752 | 0.752 |
| Symmetry imposed | C1 | C1 | C1 | C1 |
| Movies (No.) | 6,800 | 7,411 | 8,151 | 8,151 |
| Initial particle images (No.) | 1,264,111 | 903,028 | 1,370,275 | 1,370,275 |
| Final particle images (No.) | 329,210 | 387,880 | 20,011 | 87,429 |
| Map resolution (Å) | 2.1 | 2.0 | 2.6 | 2.3 |
| FSC threshold | 0.143 | 0.143 | 0.143 | 0.143 |
| Map sharpening B factor (Å$^2$) | −56.6 | −44.5 | −37.8 | −45.8 |
| 3DFSC sphericity | 0.881 | 0.886 | 0.801 | 0.854 |
| **Refinement** | | | | |
| Refinement software | Phenix | Phenix | Phenix | Phenix |
| Initial model used | AF3 | AF3 | AF3 | State III |
| Model resolution (Å) | 2.3 | 2.1 | 2.8 | 2.4 |
| FSC threshold | 0.5 | 0.5 | 0.5 | 0.5 |
| Model-Map CC (CCmask/CCbox) | 0.90/0.86 | 0.93/0.89 | 0.90/0.80 | 0.91/0.82 |
| Model composition | | | | |
| Non-hydrogen atoms | 25038 | 27007 | 34734 | 35046 |
| Protein residues | 2927 | 2926 | 3943 | 3931 |
| Nucleotide residues | 60 | 146 | 154 | 161 |
| Ligands | CPS: 5, Mg: 1 | CPS: 5, Mg: 2 | Mg: 2 | Mg: 1 |
| Average B factors (Å$^2$) | | | | |
| Protein | 126.93 | 86.90 | 97.36 | 102.71 |
| Nucleotide | 157.29 | 112.17 | 109.89 | 120.46 |
| Ligand | 111.58 | 86.11 | 121.19 | 94.56 |
| R.m.s. deviations | | | | |
| Bond lengths (Å) | 0.002 | 0.003 | 0.004 | 0.003 |
| Bond angles (°) | 0.462 | 0.455 | 0.495 | 0.460 |
| Validation | | | | |
| MolProbity score | 1.35 | 1.16 | 1.53 | 1.25 |
| Cβ outliers (%) | 0.04 | 0 | 0 | 0 |
| CaBLAM outliers (%) | 0.96 | 1.22 | 1.92 | 1.29 |
| Clashscore | 4.28 | 3.38 | 4.95 | 4.84 |
| Poor rotamers (%) | 1.48 | 1.09 | 1.42 | 0.98 |
| Ramachandran plot | | | | |
| Favored (%) | 97.93 | 98.07 | 97.11 | 98.13 |
| Allowed (%) | 2.03 | 1.93 | 2.84 | 1.87 |
| Disallowed (%) | 0.03 | 0.00 | 0.05 | 0.00 |
